# Aquatic and terrestrial organisms display contrasting life history strategies as a result of environmental adaptations

**DOI:** 10.1101/764464

**Authors:** Pol Capdevila, Maria Beger, Simone P. Blomberg, Bernat Hereu, Cristina Linares, Roberto Salguero-Gómez

## Abstract

**Aims:** Aquatic and terrestrial realms display stark differences in key environmental factors and phylogenetic composition. Despite such differences, their consequences for the evolution of species’ life history strategies remain poorly understood. Here, we examine whether and how life history strategies vary between terrestrial and aquatic species.

**Location:** Global.

**Time period:** Variable, the earliest year being in 1906 and the most recent in 2015.

**Major taxa studies:** Macroscopic animals and plants species.

**Methods:** We use demographic information for 638 terrestrial and 117 aquatic animal and plant species, to derive key life history traits capturing their population turnover, and investments in survival, development, and reproduction. We use phylogenetically corrected least squares regression to explore the differences in the trade-offs between life history traits in both realms. We then quantify the life history strategies of aquatic and terrestrial species using a phylogenetically corrected principal component analysis.

**Results:** We find that the same trade-offs structure terrestrial and aquatic life histories, resulting in two dominant axes of variation describing species’ pace- of-life and reproductive spread through time. Life history strategies differ between aquatic and terrestrial environments, with phylogenetic relationships playing a minor role. We show that adaptations of plants and animals to terrestrial environments have resulted in different life history strategies, particularly with their reproductive mode and longevity. Terrestrial plants display a great diversity of life history strategies, including the species with the longest lifespans. Aquatic animals, on the contrary, exhibit higher reproductive frequency than terrestrial animals, likely due to reproductive adaptations (i.e. internal fecundation) of the later to land.

**Main conclusions:** Our findings show that aquatic and terrestrial species are ruled by the same life history principles, but have evolved different strategies due to distinct selection pressures. Such contrasting life history strategies have important consequences for the conservation and management of aquatic and terrestrial species.

## Introduction

The rich diversity of life history strategies worldwide stem from three fundamental demographic building blocks: survival, development, and reproduction (Stearns, 1992). The life histories that emerge from the combination of these three processes determine the viability of populations (Salguero-Gómez *et al.*, 2016b; McDonald *et al.*, 2017) and guide the effectiveness of conservation plans (Paniw *et al.*, 2019). However, despite the growing body of literature in life history theory (Lande *et al.*, 2017), few studies have explicitly contrasted their validity across terrestrial and aquatic organisms (Webb, 2012).

Life history theory predicts that any strategy is shaped by two counter-acting forces: environmental filtering and phylogenetic inertia (Stearns, 1992). Regarding the former, existing environmental differences between aquatic and terrestrial realms (e.g., water density and viscosity) could have produced divergences in their respective life history strategies (Dawson & Hamner, 2008; Webb, 2012; Gearty *et al.*, 2018). Under phylogenetic inertia (Freckleton, 2000; Blomberg & Garland, 2002), life history strategies are expected to be more similar, irrespective of environment, amongst closely related lineages

Life history theory is rooted upon the concept of trade-offs as an unifying principle across the tree of life and realms (Stearns, 1992). This body of literature predicts that, due to limitations in available energy and physiological constraints, compromises among survival, development, and reproduction are inescapable (Stearns, 1992). Such constraints are expected to result in a finite set of viable demographic schedules. Indeed, comparative demographic studies have successfully identified and organised them into a few major axes of trait co-variation (Gaillard *et al.*, 1989; Salguero-Gómez *et al.*, 2016b). A seminal concept in organising such a trait co-variation is the “*fast-slow* continuum” (Stearns, 1992). In it, species are organised along two extremes: at the fast*-* living extreme, species develop fast, become highly reproductive, but die young (e.g. *Bromus tectorum* [cheatgrass], Griffith, 2010; planktonic species, Reynolds, 2006); while at the slow extreme, species develop slowly, live long, and reproduce rarely and late in life (e.g. *Somniosus microcephalus,* the Greenland shark, Nielsen *et al.*, 2016; *Pinus longaeva,* the Bristlecone pine, Peñuelas & Munné-Bosch, 2010). However, an explicit comparison of the fast-slow continuum between aquatic and terrestrial species remains, to our knowledge, untested.

Based on the strong environmental and phylogenetic differences between aquatic and terrestrial realms, life history strategies should differ between both realms. Life was originated on the sea, and the land colonisation resulted in a great divergence of the biodiversity patterns observed in both realms (Grosberg *et al.*, 2012; Costello & Chaudhary, 2017). There is a higher richness of species on land (∼80%), while aquatic biota are more diverse at the phylum level (∼34) than the terrestrial realm (∼15) (Costello & Chaudhary, 2017). Also, land colonisation required adaptations to: the effects of gravity on body structures, avoid desiccation, the elimination of waste products, together with other processes (see reviews in Grosberg *et al.*, 2012; Webb, 2012). For example, early life stages can feed and develop during dispersal in aquatic environments (Burgess *et al.*, 2016; Bush *et al.*, 2016; Vermeij & Grosberg, 2017), but terrestrial species had to evolve reproductive systems independent to environmental water, such as internal fecundity or seeds (Grosberg *et al.*, 2012; Bush *et al.*, 2016; Steele *et al.*, 2019).

The colonisation of land likely resulted in the evolution of life histories to deal with higher temporal environmental variability (Dawson & Hamner, 2008; Ruokolainen *et al.*, 2009). On land environmental variation is more random and less auto-correlated than in aquatic environments (Ruokolainen *et al.*, 2009). Classical life history theory predicts the evolution of longevity in constant environments (Lande *et al.*, 2017). However, there is recent evidence that longevity can be an strategy to deal with environmental variation (Morris *et al.*, 2008; McDonald *et al.*, 2017). On the other hand, fast life histories, are expected to show increasing fluctuations in population sizes with increasing environmental variation (Morris *et al.*, 2008; McDonald *et al.*, 2017). For that reason, some authors have argued that the colonization of land resulted in the evolution of longer lifespans to smooth out the short-term but large-amplitude terrestrial environmental fluctuations (*sensu* Steele *et al.*, 2019).

Here, we test the hypothesis that (i) trade-offs are universal both in aquatic and terrestrial systems, and (ii) that terrestrial species have evolved different life history strategies compared to aquatic ones. We use high-resolution demographic data from 117 aquatic and 638 terrestrial species across the globe from the COMPADRE and COMADRE databases (Salguero-Gómez *et al.*, 2015, 2016a). We estimate key life history traits capturing population turnover, and investments in survival, development, and reproduction of those species. To test these hypotheses, we first determine whether correlations between life history traits differ across realms as a way to examine whether trade-offs diverge between terrestrial *vs*. aquatic species. Second, we explore the main axes of life history variability shaping aquatic and terrestrial species. The presence of different life history axes of variation and/or a distinct positioning of aquatic species compared to terrestrial ones within those axes would suggest dissimilar selection pressures occurring above and below water. Given the scarcity of trans-realm comparative studies and the lack of demographic information for many aquatic species, elucidating these questions is a key step forward towards understanding the evolution of life histories across realms.

## Material and Methods

### Demographic data and life history traits

We calculated species’ life history strategies using demographic data describing information across the full life cycle of each species. This high-quality demographic information was obtained from the COMPADRE Plant Matrix Database (v. 5.0.1; Salguero-Gómez *et al.*, 2015) and COMADRE Animal Matrix Database (v. 3.0.1; Salguero-Gómez *et al.*, 2016a). In them, the demographic data is archived into matrix population models (MPMs, hereafter) for over 700 plant and 400 animal species, respectively. MPMs are summaries of organisms’ demographic processes (*i.e.,* vital rates) that together determine their life history strategies and resulting population dynamics (Caswell, 2001). For this reason, MPMs provide the ideal means to compare the vast array of life history strategies (Franco & Silvertown, 2004; McDonald *et al.*, 2017).

To compare life history traits across aquatic and terrestrial species, we imposed a series of selection criteria to the available demographic data (see details in Appendix S2: Data selection in Supporting information). These criteria resulted in 638 terrestrial species and 117 aquatic species used in this study (Appendix S1). We also classified aquatic vs terrestrial species according to the information provided in the World’s Register of Marine Species (http://www.marinespecies.org) and the Catalogue of Life (http://www.catalogueoflife.org). The number of species studied here represented a similar taxonomic coverage relative to the known biodiversity of the aquatic (∼0.05%) and terrestrial realm (∼0.01%; Table S1 in Appendix S2).

Quantifying a species’ life history strategy requires detailed information regarding the timing, intensity, frequency, and duration of key demographic processes across its life cycle (Stearns, 1992). To quantify species’ life history strategies, we calculated several life history traits from each MPM that are *a priori* not correlated using well-established methods (Salguero-Gómez *et al.*, 2016b). We selected seven life history traits commonly used in comparative demography (Stearns, 1992; Gaillard *et al.*, 2005; Bielby *et al.*, 2007; Salguero-Gómez *et al.*, 2016). These traits include: generation time (*T*), age at sexual maturity (*L*_α_), rate of senescence (*H*), mean vital rate of progressive development (*γ*), the mean vital rate of sexual reproduction (*ϕ*) and degree of iteroparity (*S*) (Table S2). Such traits provide insights of the species’ population turnover, as well as of survival, developmental, and reproductive strategies (detailed in Table S2 in Appendix S2).

### Phylogenetic analyses and trait comparisons

We accounted for and estimated the phylogenetic influence on the differences in life history trait values within species and between aquatic *vs.* terrestrial realms. To do so, we constructed a species-level phylogenetic tree (Figure S2 in Appendix S3) with data from Open Tree of Life (OTL, https://tree.opentreeoflife.org, Hinchliff *et al.*, 2015). OTL combines publicly available taxonomic and phylogenetic information across the tree of life (Hinchliff *et al.*, 2015). Briefly, we built separate trees for our species of algae, plant, and animals, using the *rotl* R package (Michonneau *et al.*, 2016), which were assembled in a supertree using the function *bind.tree* in the *phytools* package (Revell, 2012). To account for the phylogenetic relatedness of species we computed the branch lengths and resolved polytomies (Revell, 2012). We also tested the sensitivity of our results to the choice of a particular set of branch lengths, by repeating our analyses setting all the branch lengths to one and using Pagel’s branch length (Tables S4-S7 in Appendix S3) using the software Mesquite 1.05 (Maddison & Maddison, 2001) and its PDAP module 1.06 (Midford *et al.*, 2005). For further details on the construction of the tree see Appendix S3.

To test whether life history trait trade-offs are congruent between aquatic *vs.* terrestrial species, we carried out a series of Phylogenetic General Least Square (PGLS) analyses (Revell, 2010). This approach allows us to accommodate residual errors according to a variance-covariance matrix that includes ancestral relationships between any pair of species from our phylogenetic tree (Revell, 2010, 2012). We implemented our set of PGLSs in R using the correlation structures provided by the package *ape* (Paradis *et al.*, 2004). We used a Brownian motion model of evolution (BM), combined with the pgls function from the *nlme* package (Pinheiro *et al.*, 2014). Separate PGLSs were fitted using Ornstein Uhlenbeck (OU) model of evolution, which describes Brownian model under the influence of friction (Uhlenbeck & Ornstein, 1930). Both models where compared using Akaike Information Criterion (Akaike, 1974); the BM generally outperformed the OU model, but both showed similar results. Therefore, we only report the PGLS results from the Brownian motion model.

### Exploring dominant axes of life history strategies

To explore the patterns of association among life history traits for aquatic *vs*. terrestrial species, we performed a series of principal components analyses (PCA). PCA is a multivariate analysis that reduces a set of correlated variables into linearly uncorrelated measurements, the so-called principal components (PCs). Life history trait data were log- and z-transformed (mean=0, SD=1) to fulfil normality assumptions of PCAs (Legendre & Legendre, 2012). Finally, we identified and excluded outliers for each life history trait as those located outside of the 2.5^th^-97.5^th^ percentile range of the distribution. However, we note that the exclusion of outliers did not alter our main findings (see Tables S8-S11 in Appendix S4).

To account for shared ancestry while exploring differences in aquatic *vs.* terrestrial life history strategies, we used a phylogenetically informed PCA (pPCA Revell, 2009). The pPCA considers the correlation matrix of species’ traits while accounting for phylogenetic relationships and simultaneously estimating Pagel’s *λ* with maximum likelihood methods. Pagel’s *λ* quantifies the strength of the phylogenetic relationships on trait (co-)evolution under a BM (Freckleton, 2000; Blomberg & Garland, 2002). This metric varies between 0 when the observed patterns are not due to phylogenetic relationships, and 1 when the observed patterns can be explained by the employed phylogeny (Blomberg & Garland, 2002; Revell, 2010). The pPCA was estimated using the phyl.pca function from the R package *phytools* (Revell, 2012), assuming a BM (Revell, 2010).

A full dataset (*i.e.*, no missing values) is necessary to run the pPCA analyses. However, estimating life history traits for species’ MPMs was not always possible (see *Missing data* in Appendix S2: Extended methods). For example, we could not calculate the rate of senescence for *Fucus vesiculosus*. The rate of senescence (Keyfitz’s entropy) can only be calculated for life tables that have not reached stationary equilibrium before the 95% of a cohort are dead (Caswell, 2001; Jones *et al.*, 2014), which was not the case for this species. In these cases, we imputed the missing data using function *amelia* from the *Amelia* package (Honaker *et al.*, 2011). This function uses a bootstrap EM algorithm to impute missing data. We then created 10 imputed datasets and ran analyses on each separately. In addition, we tested the sensitivity of our results to missing traits in the dataset using pPCA in two ways. First, we ran a pPCA only with species without any missing data (62 aquatic species, 477 terrestrial species, Tables S10 and S11 in Appendix S4), and with missing species trait values filled using imputation methods (see Tables S8, S9, S12 and S13 in Appendix S4). The results from the multiple imputations were presented as their respective mean values with their standard deviation. To test the differences between the distributions of pPCA scores between realms, we used the mean position resulting from the multiple imputations.

We also examined the consistency of our results and explored the differences between realms by performing the pPCA analyses on different subsets of data. These subsets included comparisons between mobile *vs.* sessile organisms, Animalia *vs.* Plantae/Chromista kingdoms, and aquatic *vs.* terrestrial realms. We considered sessile species as those that do not have active locomotion during the adult stages of their life cycle (e.g. corals, sponges, plants) and those species with limited adult locomotion (e.g. clams, worms, snails). This distinction was made because key traits (e.g. reproduction, development, energetic requirements) can differ between sessile and mobile organisms (Bush *et al.*, 2016; Vermeij & Grosberg, 2017). We also performed a series of pPCA analyses sub-setting species into Animalia kingdom, and Plantae and Chromista (brown algae). This distinction was also made because animals and plants/algae differ in key physiological, trophic and development traits (Grosberg *et al.*, 2012; Burgess *et al.*, 2016). Such ecological differences between sessile/mobile and taxonomic kingdoms, could have a potential impact on our hypothesis about the different evolution of life history strategies in aquatic and terrestrial species.

## Results

### Trade-offs are pervasive across realms

Life history traits are shaped by the same trade-offs below water as on land (Figure 1). Our PGLS analyses reveal a similar magnitude and the same direction of pair-wise correlations between traits for aquatic and for terrestrial species (Figure 1 and Tables S9, S11 and S13 in Appendix S4). Regardless of the realm, producing many recruits (high *φ*; Table S2 in Appendix S2) results in fast population turnover (low *T*). Likewise, species that postpone their first reproductive event (high *L*_*α*_) have low senescence rates (high *H*) (Figure 1). Species with fast development (high *γ*) achieve reproductive maturity early (low *L*_*α*_) at the cost of high senesce rates (low *H*). Also, those species high reproductive output (high *ϕ*) and frequent reproduction (high *S*), have low senescence rates (high *H*) (Figure 1).

**Figure 1.**
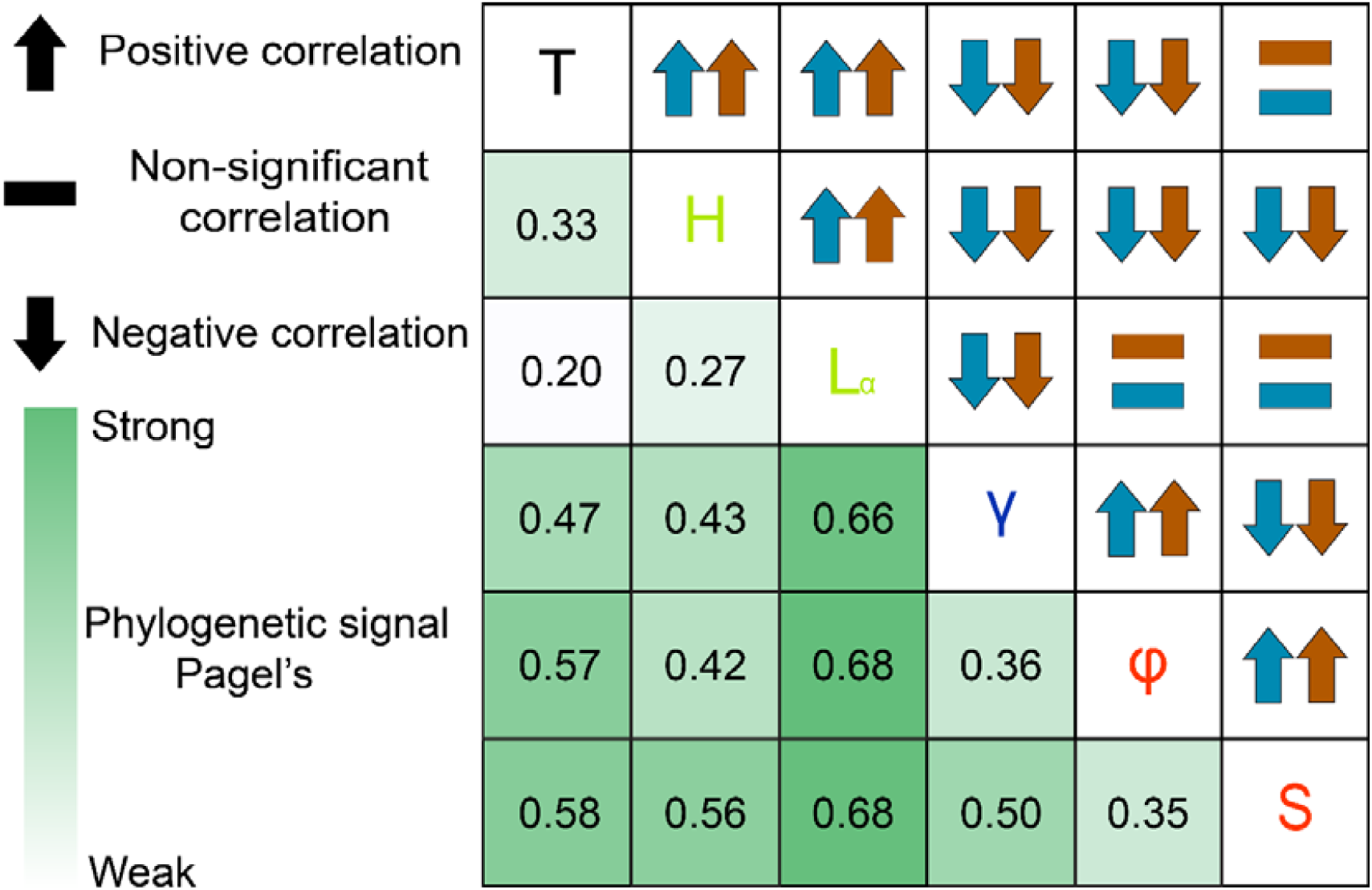
Trade-offs among life history traits are congruent between aquatic and terrestrial environments. Pair-wise correlations between seven life history traits (Table S4) for 117 aquatic (blue) and 638 terrestrial (brown) species. Arrows indicate the direction of each pair-wise correlation using phylogenetic generalised least squares: positive (arrow-up), negative (arrow-down) or not-significant correlation (horizontal bar; *P*>0.05). The mean phylogenetic signal (Pagel’s *λ*) of each pair-wise correlation, displayed in the lower-triangle, ranges from weak (white, ∼0.1) to strong (dark green, ∼0.9).

### Aquatic species are faster than terrestrial ones

Together, the first two axes of our phylogenetically corrected principal component analysis (pPCA; Table 1) explain over 60% of the examined variation in life history traits (Figure 2, Table 1). Principal component axis 1 (PC1) explains 47.42±0.34% (Mean±S.E.) of the variation and represents the fast*-*slow continuum. Indeed, PC1 portrays a trade-off between species with fast development and short lifespans, and slow development, high investment in survival (low senescence rates), and postponement of maturity (Figure 2). PC2 explains 21.02±0.11% of the variation in life history traits related to reproductive strategies. In PC2, those species characterised by high reproductive rate and high iteroparity are located at the top *vs.* species with fewer reproductive events across their lifetimes, located at the bottom of Figure 2. These patterns are robust within different life modes (Figure 3a,b and Table S14 in Appendix S4), kingdoms (Figure 3c,d and Table S15 in Appendix S4), and realms (Table S16 in Appendix S4).

**Table 1.**
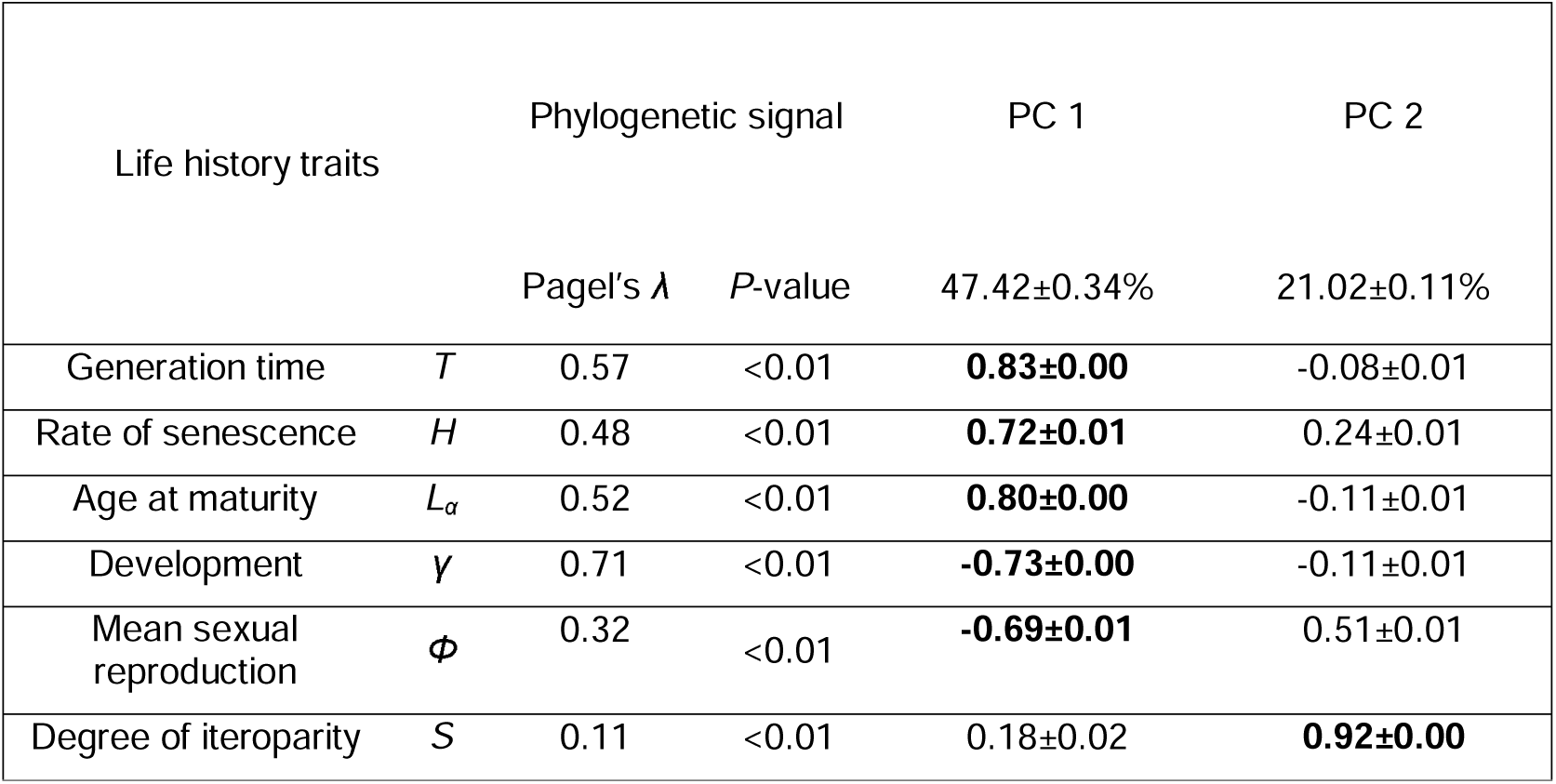
Life history traits used in the comparative analyses of 638 terrestrial and 117 aquatic species to examine differences in strategies between both realms, together with their loadings on the first two principal component axes, grouped by their attribution to turnover, survival, development, or reproduction. Pagel’s *λ* (and its associated *P*-value) describes the strength of phylogenetic inertia, ranging between 1, when life history trait differences are entirely due to the phylogenetic structure of the data under Brownian motion, and 0, meaning no phylogenetic structuring in the pattern. The mean loading values of each life history trait are visually depicted in Figure 2A. SE values were calculated via 10 imputations (See Methods). Bold numbers indicate traits loadings above 55% for each PC.

**Figure 2.**
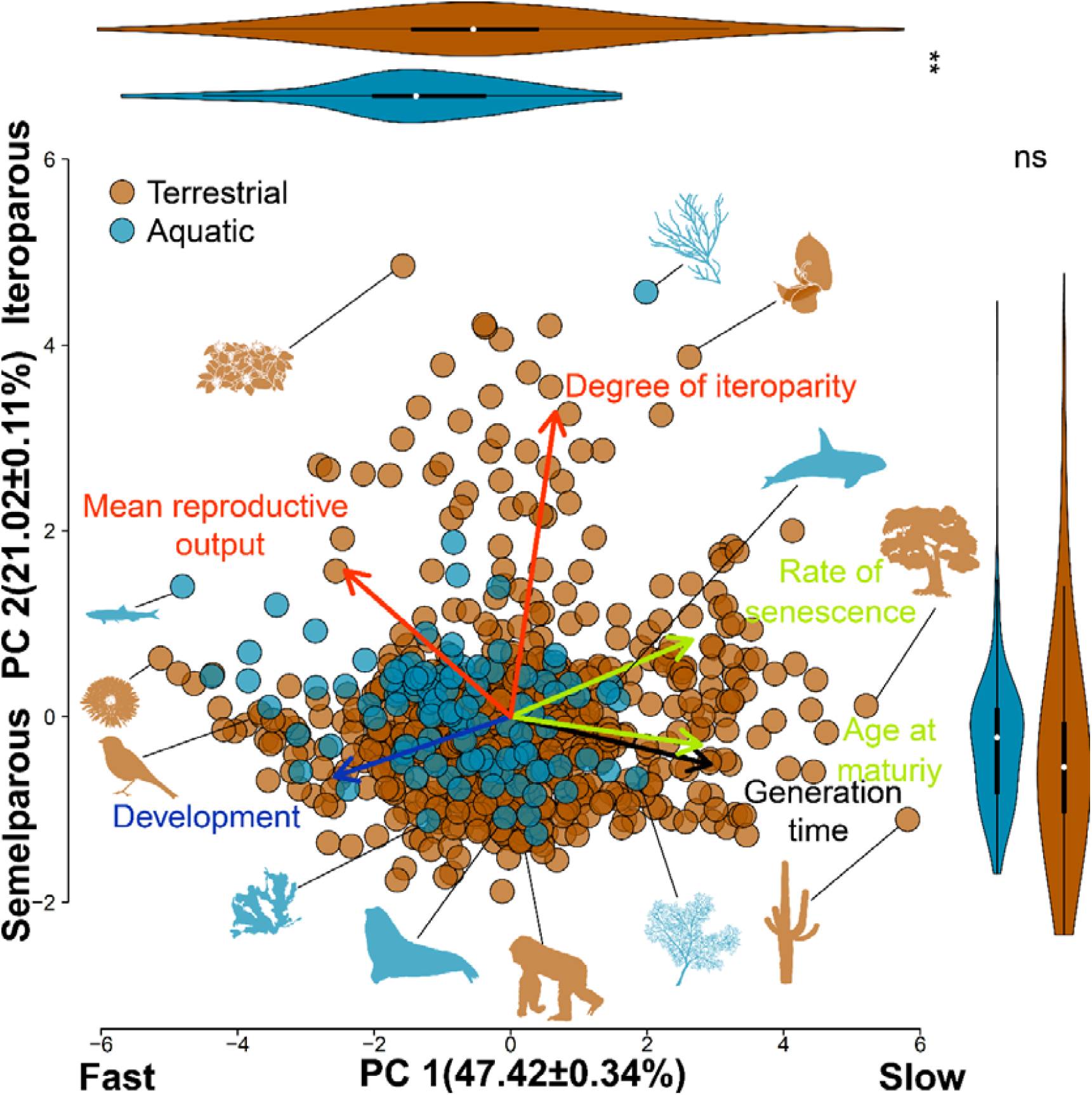
Aquatic and terrestrial life history strategies are organised in two main axes of variation, the fast-slow continuum and the reproductive strategies. Phylogenetically-corrected principal component analysis (pPCA) for the first two axes (percentage of variance absorbed in brackets) for seven key life history traits from 117 aquatic (blue) and 638 terrestrial species (brown). Arrow lengths indicate mean loading of each life history trait, and colour indicates associations with population turn-over (black), survival (green), development (dark blue), and reproduction (red). Each point represents the mean position of a species on this two-dimensional space for 10 imputed data sets (see Methods). Violin plots (top and right) depict the distribution of species along each PC axis; white dot: mean; black thick line: 25^th^-75^th^ quantile; black thin line: SD; ns: not-significant; *: *P*<0.01; **: *P*<0.005. The silhouettes, starting at the top left, and counter-clock-wise, correspond to: *Lantana camara, Clinostomus funduloides, Solidago mollis, Setophaga cerulea, Mazzaella splendens, Mirounga leonina, Gorilla beringei, Paramuricea clavata, Pseudomitrocereus fulviceps, Chlorocardium rodiei, Orcinus orca, Cypripedium calceolus* and *Gracilaria gracilis.*

**Figure 3.**
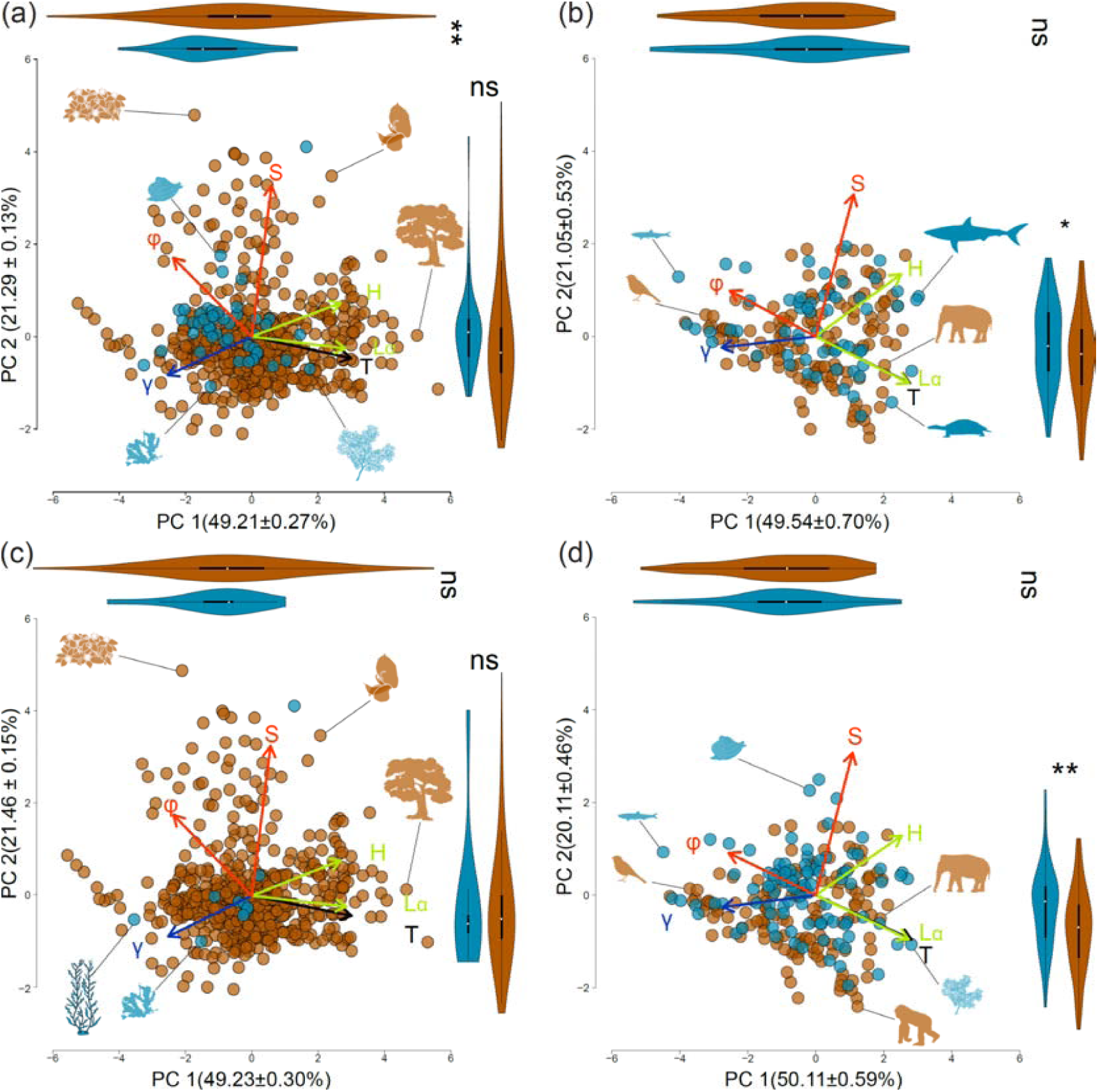
The main axes of life history variation remain constant, regardless of the degree mobility/sessility or taxonomic kingdom. Phylogenetically-corrected principal component analysis of seven life history traits across 683 species. Trait definition in Figure 2 and Table S4. Note that the fast-slow continuum remains the dominant axis of variation, explaining 40-45% of the variation, followed by an axis of reproductive strategies, which explains 20-22% of the variation in life history traits. ns: not-significant; *: *P*<0.05; **: *P*<0.01. (a) Sessile organisms, with silhouettes (not to scale; starting at the top left, and counter-clock-wise) representing: *Lantana camara, Mya arenaria, Mazzalena splendens, Paramuricea clavata, Chlorocardium rodiei* and *Cypripedium calceolus*. (b) Mobile organisms: *Clinostomus funduloides, Setophaga cerulea, Chelydra serpentine, Elephas maximus* and *Isurus oxyrhincus* (c) Kingdoms Plantae and Chromista: *L. camara, Pterygophora caliphornica, M. splendens C. rodiei* and *C. calceolus*. (d) Kingdom Animalia: *M. arenaria, C. funduloides, S. cerulea, Gorilla beringei, P. clavata* and *E. maximus*.

Aquatic life history strategies are displaced towards the fast extreme of the fast*-*slow continuum (*t*_197.49,_=-6.22, *P*<0.01; Figure 2). On land, the studied species occupy fast paces of life, such as *Solidago mollis* (soft goldenrod), *Setophaga cerulean* (cerulean warbler), as well as slow ones, such as *Pseudomitrocereus fulviceps* (the giant cardon). In the aquatic realm, in contrast, the resulting paces of life are constrained to faster values compared to terrestrial species (PC1, Figure 2). In contrast, aquatic organisms are not displaced towards any of the extremes of the PC2. Both aquatic and terrestrial species show a wide range of reproductive strategies, with highly reproductive species such as such as *Lantana camara* (big-sage) or *Gracilaria gracilis* (red seaweed), and species less reproductive species, such as *Mirounga leonina* (southern elephant seal) and *Gorilla beringei* (eastern gorilla). There are no significant differences between the realms in the PC2 (*t*_215.04_= 0.18, *P*=0.86; Figure 2).

### Mode-of-life and kingdom drive key life history differences across realms

The main axes of life history variation remain unaltered across realms, modes-of-life (whether species are mobile or sessile during their adulthood) or taxonomic affiliation. The first and second axes of life history trait variation correspond to the fast*-*slow continuum and reproductive strategies in both sessile and mobile species (Figure 3a,b and Table S14 in Appendix S4), in Animalia and Plantae/Chromista kingdoms (Figure 3c,d and Table S15 in Appendix S4), in terrestrial species and aquatic species (Table S16 in Appendix S4).

Aquatic and terrestrial sessile species display significant differences in their position across first axes of life history variation. Aquatic sessile species are displaced towards the fast end (*i.e.,* low PC1 scores) of the fast-slow continuum (*t*_64.91_=-53.32, *P*<0.01; Figure 3a). Aquatic sessile species do not show significant differences in their reproductive strategies compared to terrestrial ones (*t*_59.22_=1.95, *P*=0.06; Figure 3a). In contrast, mobile aquatic species are not displaced towards any end of the fast-slow continuum when compared to terrestrial mobile species (*t*_96.34_=0.55, *P*=0.58; Figure 3b), neither in the reproductive axis (*t*_118.88_=1.84, *P*=0.07; Figure 3b).

Terrestrial plants have a wide range of life history strategies with no significant displacement in the pace-of-life axis (*t*_9.52_=-1.16, *P*=0.27; Figure 3c) neither in the reproductive strategies’ axis (*t*_9.16_=0.25, *P*=0.81; Figure 3c). In contrast, animals do not show any significantly displaced towards any end of the fast-slow continuum (*t*_199.08_=0.74, *P*=0.46; Figure 3d). In contrast, aquatic animals are significantly displaced towards the upper end of the reproductive axis compared to their terrestrial counterparts (*t*_208.60_= 4.27, *P*<0.01; Figure 3d).

### Ancestry does not shape cross-realm life history strategies

Overall, phylogenetic ancestry (*i.e.,* phylogenetic inertia) plays a minor role in constraining life history strategies between realms. The estimates of Pagel’s *λ* in our pPCA are indeed low (0.26±0.00). Such values of the phylogenetic signal remain low across sessile (*λ*=0.18±0.01; Table S14 in Appendix S4), mobile species (*λ*=0.36±0.01; Table S14 in Appendix S4), plants and algae (*λ*=0.18±0.01; Table S15 in Appendix S4) and animals (*λ*=0.31±0.02; Table S15 in Appendix S4). In addition, the phylogenetic signal is similar between terrestrial (*λ*=0.24±0.01; Table S16 in Appendix S4) and aquatic species (*λ*=0.19±0.02; Table S16 in Appendix S4).

For both aquatic and terrestrial species, reproductive traits (□ and S in Table S2) are systematically more labile (*i.e.*, lower phylogenetic signal) than traits associated to survival (*H, L*_*α*_), development (*γ*) or turnover (*T*). Generation time (*T*) and age at reproductive maturity (*L*_*α*_) are strongly phylogenetically associated with the number of recruits produced (□) and the degree of iteroparity (*S*)(Fig.1). The traits with the highest loading on the fast-slow continuum (*T, H* and *L*_*α*_) are strongly phylogenetically linked to two leading traits of the reproductive-strategies axis (□ and S). Equally, the variation on age at maturity (*L*_*α*_) is largely explained by its phylogenetic association with developmental rates (*γ*) (Figure 1).

## Discussion

Our comparison of 117 aquatic and 638 terrestrial species demonstrates that life history strategies are organised along the same dominant axes of variation and constrained by the same trade-offs, regardless of realm. The sampled aquatic species have not evolved the high longevities attained by some of our studied terrestrial species, but those aquatic animals are more reproductive than the terrestrial ones. The relatively weak phylogenetic signal in our analyses suggest that these key life history differences are not explained by the differential taxonomic composition of both realms (Freckleton, 2000; Blomberg & Garland, 2002). Overall, we suggest that the contrasting environmental conditions between aquatic and terrestrial realms may play a major role in the observed life history patterns and differences.

We show greater diversity of life history strategies on land compared to aquatic environments. This finding is congruent with the higher species richness (Costello & Chaudhary, 2017) and larger range of species biomass housed in the terrestrial realm (Bar-On *et al.*, 2018). The colonisation of land established a period of unparalleled innovations in the evolution of plants and animals, driven by challenges in water retention, mobility, and dispersal (Steele *et al.*, 2019). Adaptations like plant vascularity, and animal terrestrial mobility were key for the proliferation of populations and species diversification (Steele *et al.*, 2019). These innovations allowed the exploitation of novel ecological niches, ultimately resulting in a six-fold increase in speciation rate (Costello & Chaudhary, 2017). We argue that such adaptations are reflected in the vast diversity of life histories observed in the terrestrial realm relative to that in the aquatic realm in our study.

Nevertheless, plants and animals evolved different sets of adaptations to terrestrial and aquatic environments (Burgess *et al.*, 2016; Steele *et al.*, 2019), resulting in distinct life history strategies too. Terrestrial plants account for most of the diversity of life histories observed in our study, but they show slower strategies than aquatic species. Slow life history strategies can buffer environmental variation compensating the uncertainties of reproductive success through high adult survival (Morris *et al.*, 2008; McDonald *et al.*, 2017), and have been suggested as an adaptation of plants to terrestrial environments (Steele *et al.*, 2019). Such a pattern, however, is not shared with terrestrial animals, which do not show any displacement towards any extreme of the fast-slow continuum, compared to aquatic animals. In contrast, terrestrial animals could have compensated environmental uncertainties through the evolution of complex behaviours (e.g. societies, nesting) and physiological adaptations (e.g. thermoregulation, internal fecundation) (Grosberg *et al.*, 2012; Steele *et al.*, 2019).

A major challenge for land colonisation is reproducing in a non-aquatic environment (Grosberg *et al.*, 2012; Burgess *et al.*, 2016), influencing the evolution of divergent life history strategies. External fertilisation is the predominant reproductive strategy in aquatic environments (Bush *et al.*, 2016). Aquatic colonisers to land environments had to evolve strategies to protect early stages (e.g. to desiccation) and enable their development in non-aquatic environments (Strathmann, 1990; Burgess *et al.*, 2016; Steele *et al.*, 2019). Plants, like many benthic aquatic species, have a sessile adulthood, so their dispersal is in early developmental stages only. This life-mode promoted the evolution of flowers, pollination and seeds (Kenrick & Crane, 1997), resulting in the observed high reproductive outputs and frequencies in plants, despite the fact that they can also reach high longevities (e.g. Salguero-Gómez *et al.*, 2016b; McDonald *et al.*, 2017).

On the other hand, the colonisation of land constrained animal reproduction to internal fertilisation, with a consequent decrease in reproductive outputs and frequency. Again, we argue that this finding is likely linked to the prevalence of external fertilisation in the ocean compared to the terrestrial realm (Bush *et al.*, 2016). Both viscosity and nutrient concentration are higher in seawater than in air (Dawson & Hamner, 2008), allowing propagules to remain suspended for long periods of time (Strathmann, 1990; Burgess *et al.*, 2016). The release of progeny in the water column comes with a high early predation risk and low establishment probability and high early mortality (Strathmann, 1990; Burgess *et al.*, 2016). To compensate such early mortality, aquatic species release high numbers of propagules and frequently, resulting in highly reproductive life histories. In contrast, most terrestrial animals retain female gametes on or in their bodies, and fertilisation and early development are usually internal (Bush *et al.*, 2016; Steele *et al.*, 2019), resulting in less reproductive strategies. Still, there are aquatic species that have internal fecundity, such as some sharks or marine mammals (Steele *et al.*, 2019), explaining the range of reproductive strategies observed in our study.

Although the volume of data used in our study has a similar ratio to that of the biodiversity held in aquatic *vs.* terrestrial realms (∼0.01%; Table S1 in Appendix S1), it still represents a limited fraction of the known diversity (Grosberg *et al.*, 2012; Costello & Chaudhary, 2017). Importantly, here we have focused mostly on macroscopic organisms, for which full demographic information is more readily available than for small species (Salguero-Gómez *et al.*, 2015, 2016a). Organisms like insects, but also microscopic organisms, such as plankton or bacteria, are challenging subjects for demographic studies, so their data are very scarce (Salguero-Gómez *et al.*, 2016a; Conde *et al.*, 2019). In addition, recently discovered extremely long-lived marine species (e.g. *Somniosus microcephalus*, Nielsen *et al.*, 2016; *Monorhaphis chuni*, Jochum *et al.*, 2012) are likely examples of slow strategies for which we do not yet have complete demographic data. Thus, the increase of studies quantifying the demographic processes of the full life cycle of species will likely shed more light on the differences between aquatic and terrestrial life histories.

In this study, we used demography as the common currency to quantify the life history strategies of species. Species life history strategies are highly determined by the demographic processes of survival, development and reproduction (Stearns, 1992; Caswell, 2001). Researchers quantifying life history strategies have used different approaches to compare species (e.g. fishes in Winemiller & Rose, 1992; plants in Westoby, 1998; Grime & Pierce, 2012). These approaches have significantly contributed to improve our current understanding of life history strategies both in terrestrial and aquatic realms (Grime & Pierce, 2012). However, in some cases, these approaches use taxon-specific traits (such as the leaf–height–seed strategy scheme by Westoby, 1998), which would not allow us to compare across different taxonomic groups, such as animals and plants. For that reason, demographic data quantifying important moments of the life cycle of species (Salguero-Gómez *et al.*, 2016b), provides the ideal means to compare strategies across very different and distant taxonomic groups. Importantly, we also demonstrate that considering incomplete demographic information (e.g. only survival investments) can lead to the inaccurate characterisation of the life history strategy of a given species. Information on the pace-of-life explains the life history variation of about 52.52% of aquatic and 49.05% of terrestrial species. Typically lacking reproductive information, which is much more challenging to collect than to estimate survival, prevents us from improving our understanding by 21.61% and 21.35%, of aquatic and terrestrial species respectively.

Overall, our study provides a promising entry-point to trans-realm comparative biology. Our findings evidence the existence of strong differences between the life history strategies of aquatic and terrestrial systems as a consequence of the colonization of land environments. Such contrasting life history strategies are probably linked to the distinct responses to climate change (Pinsky *et al.*, 2019), exploitation (McCauley *et al.*, 2015) or extinction rates (Webb & Mindel, 2015) observed in aquatic and terrestrial systems. Understanding how the contrasting patterns of life histories translate into differences in their response to disturbances will be crucial to improve management decisions and predict future biodiversity trends.

## Acknowledgements

We thank T. Coulson, H. Possingham, O. Jones, and F. Colchero for feedback to early versions of the manuscript, and the Max Planck Institute for Demographic Research for the development of and access to the COMPADRE and COMADRE databases. P.C. was supported by a FI-DRG grant from the Generalitat de Catalunya, the Smart project (CGL2012-32194) funded by the Spanish Ministry of Economy and Innovation and by a Ramón Areces Foundation Postdoctoral Scholarship. M.B. was supported by ARC CE110001014, a University Academic Fellowship by the University of Leeds, and EU MSCA DLV-747102. This research emerged through funding by ARC DE140100505 and NERC R/142195-11-1 to R.S-G.

## Data accessibility statement

Matrix population models are available at http://www.compadre-db.org.

## References

Akaike, H. (1974) A new look at the statistical model identification. IEEE Transactions on Automatic Control, 19, 716–723.

Bar-On, Y.M., Phillips, R. & Milo, R. (2018) The biomass distribution on Earth. Proceedings of the National Academy of Sciences, 201711842.

Blomberg, S.P. & Garland, T. (2002) Tempo and mode in evolutionl□: phylogenetic inertia, adaptation and comparative methods. Journal of Evolutionary Biology, 15, 899–910.

Burgess, S.C., Baskett, M.L., Grosberg, R.K., Morgan, S.G. & Strathmann, R.R. (2016) When is dispersal for dispersal? Unifying marine and terrestrial perspectives. Biological Reviews, 91, 867–882.

Bush, A.M., Hunt, G. & Bambach, R.K. (2016) Sex and the shifting biodiversity dynamics of marine animals in deep time. Proceedings of the National Academy of Sciences, 113, 14073–14078.

Caswell, H. (2001) Matrix Population Models: Construction, Analysis, and Interpretation, 2nd edn. Sinauer Associates.

Conde, D.A., Staerk, J., Colchero, F., Silva, R. da, Schöley, J., Baden, H.M., Jouvet, L., Fa, J.E., Syed, H., Jongejans, E., Meiri, S., Gaillard, J.-M., Chamberlain, S., Wilcken, J., Jones, O.R., Dahlgren, J.P., Steiner, U.K., Bland, L.M., Gomez-Mestre, I., Lebreton, J.-D., Vargas, J.G., Flesness, N., Canudas-Romo, V., Salguero-Gómez, R., Byers, O., Berg, T.B., Scheuerlein, A., Devillard, S., Schigel, D.S., Ryder, O.A., Possingham, H.P., Baudisch, A. & Vaupel, J.W. (2019) Data gaps and opportunities for comparative and conservation biology. Proceedings of the National Academy of Sciences, 116, 9658–9664.

Costello, M.J. & Chaudhary, C. (2017) Marine biodiversity, biogeography, deep-sea gradients, and conservation. Current Biology, 27, R511–R527.

Dawson, M.N. & Hamner, W.M. (2008) A biophysical perspective on dispersal and the geography of evolution in marine and terrestrial systems. Journal of the Royal Society, Interface / the Royal Society, 5, 135–150.

Franco, M. & Silvertown, J. (2004) A comparative demography of plants based upon elasticities of vital rates. Ecology, 85, 531–538.

Freckleton, R.P. (2000) Phylogenetic tests of ecological and evolutionary hypotheses: Checking for phylogenetic independence. Functional Ecology, 14, 129–134.

Gaillard, J.-M., Pontier, D., Allainé, D., Lebreton, J.D., Trouvilliez, J. & Clobert, J. (1989) An analysis of demographic tactics in birds and mammals. Oikos, 56, 59–76.

Gearty, W., McClain, C.R. & Payne, J.L. (2018) Energetic tradeoffs control the size distribution of aquatic mammals. Proceedings of the National Academy of Sciences, 115, 4194–4199.

Griffith, A.B. (2010) Positive effects of native shrubs on *Bromus tectorum* demography. Ecology, 91, 141–154.

Grime, J.P. & Pierce, S. (2012) The evolutionary strategies that shape ecosystems, John Wiley & Sons.

Grosberg, R.K., Vermeij, G.J. & Wainwright, P.C. (2012) Biodiversity in water and on land. Current Biology, 22, R900–R903.

Hinchliff, C.E., Smith, S.A., Allman, J.F., Burleigh, J.G., Chaudhary, R., Coghill, L.M., Crandall, K.A., Deng, J., Drew, B.T., Gazis, R., Gude, K., Hibbett, D.S., Katz, L.A., Laughinghouse, H.D., McTavish, E.J., Midford, P.E., Owen, C.L., Ree, R.H., Rees, J.A., Soltis, D.E., Williams, T. & Cranston, K.A. (2015) Synthesis of phylogeny and taxonomy into a comprehensive tree of life. Proceedings of the National Academy of Sciences, 112, 12764–12769.

Honaker, J., King, G. & Blackwell, M. (2011) Amelia II: A program for missing data. Journal of Statistical Software, 45, 1–2.

Jochum, K.P., Wang, X., Vennemann, T.W., Sinha, B. & Müller, W.E.G. (2012) Siliceous deep-sea sponge *Monorhaphis chuni*: A potential paleoclimate archive in ancient animals. Chemical Geology, 300–301, 143–151.

Jones, O.R., Scheuerlein, A., Salguero-Gómez, R., Camarda, C.G., Schaible, R., Casper, B.B., Dahlgren, J.P., Ehrlén, J., García, M.B., Menges, E.S., Quintana-Ascencio, P.F., Caswell, H., Baudisch, A. & Vaupel, J.W. (2014) Diversity of ageing across the tree of life. Nature, 505, 169–73.

Kenrick, P. & Crane, P.R. (1997) The origin and early evolution of plants on land. Nature, 389, 33–39.

Lande, R., Engen, S. & Sæther, B.-E. (2017) Evolution of stochastic demography with life history tradeoffs in density-dependent age-structured populations. Proceedings of the National Academy of Sciences, 114, 11582–11590.

Legendre, P. & Legendre, L. (2012) Numerical Ecology, 3rd edn. Elsevier Science, Amsterdam.

Maddison, W.P. & Maddison, D.R. (2001) Mesquite: a modular system for evolutionary analysis.

McCauley, D.J., Pinsky, M.L., Palumbi, S.R., Estes, J. a., Joyce, F.H. & Warner, R.R. (2015) Marine defaunation: Animal loss in the global ocean. Science, 347, 247–254.

McDonald, J.L., Franco, M., Townley, S., Ezard, T.H.G., Jelbert, K. & Hodgson, D.J. (2017) Divergent demographic strategies of plants in variable environments. Nature Ecology and Evolution, 1, 29.

Michonneau, F., Brown, J. & Winter, D. (2016) rotl: An R package to interact with the Open Tree of Life data. Methods in Ecology and Evolution, 7, 1–17.

Midford, P., Garland Jr, T. & Maddison, W. (2005) PDAP package of Mesquite.

Morris, W.F., Pfister, C.A., Tuljapurkar, S., Haridas, C.V., Boggs, C.L., Boyce, M.S., Bruna, E.M., Church, D.R., Coulson, T., Doak, D.F., Forsyth, S., Gaillard, J.-M., Horvitz, C.C., Kalisz, S., Kendall, B.E., Knight, T.M., Lee, C.T. & Menges, E.S. (2008) Longevity Can Buffer Plant and Animal Populations Against Changing Climatic Variability. Ecology, 89, 19–25.

Nielsen, J., Hedeholm, R.B., Heinemeier, J., Bushnell, P.G., Christiansen, J.S., Olsen, J., Ramsey, C.B., Brill, R.W., Simon, M., Steffensen, K.F. & Steffensen, J.F. (2016) Eye lens radiocarbon reveals centuries of longevity in the Greenland shark (*Somniosus microcephalus*). Science, 353, 702–704.

Paniw, M., Maag, N., Cozzi, G., Clutton-Brock, T. & Ozgul, A. (2019) Life history responses of meerkats to seasonal changes in extreme environments. Science, 363, 631–635.

Paradis, E., Claude, J. & Strimmer, K. (2004) APE: Analyses of phylogenetics and evolution in R language. Bioinformatics, 20, 289–290.

Peñuelas, J. & Munné-Bosch, S. (2010) Potentially immortal? New Phytologist, 187, 564–567.

Pinheiro, J., Bates, D., Debroy, S., Sarkar D & R Core Team (2014) nlme: Linear and nonlinear mixed effects models. R package version 3.1-117.

Pinsky, M.L., Eikeset, A.M., McCauley, D.J., Payne, J.L. & Sunday, J.M. (2019) Greater vulnerability to warming of marine versus terrestrial ectotherms. Nature, 569, 108.

R Core Team (2015) R: a language and environment for statistical computing.

Revell, L.J. (2010) Phylogenetic signal and linear regression on species data. Methods in Ecology and Evolution, 1, 319–329.

Revell, L.J. (2012) phytools: An R package for phylogenetic comparative biology (and other things). Methods in Ecology and Evolution, 3, 217–223.

Revell, L.J. (2009) Size-correction and principal components for interspecific comparative studies. Evolution, 63, 3258–3268.

Reynolds, C. (2006) Ecology of Phytoplankton, Cambridge University Press, Cambridge.

Ruokolainen, L., Lindén, A., Kaitala, V. & Fowler, M.S. (2009) Ecological and evolutionary dynamics under coloured environmental variation. Trends in Ecology & Evolution, 24, 555–563.

Salguero-Gómez, R., Jones, O.R., Archer, C.R., Bein, C., de Buhr, H., Farack, C., Gottschalk, F., Hartmann, A., Henning, A., Hoppe, G., Römer, G., Ruoff, T., Sommer, V., Wille, J., Voigt, J., Zeh, S., Vieregg, D., Buckley, Y.M., Che-Castaldo, J., Hodgson, D., Scheuerlein, A., Caswell, H. & Vaupel, J.W. (2016a) COMADRE: A global data base of animal demography. Journal of Animal Ecology, 85, 371–384.

Salguero-Gómez, R., Jones, O.R., Archer, C.R., Buckley, Y.M., Che-Castaldo, J., Caswell, H., Hodgson, D., Scheuerlein, A., Conde, D.A., Brinks, E., de Buhr, H., Farack, C., Gottschalk, F., Hartmann, A., Henning, A., Hoppe, G., Römer, G., Runge, J., Ruoff, T., Wille, J., Zeh, S., Davison, R., Vieregg, D., Baudisch, A., Altwegg, R., Colchero, F., Dong, M., de Kroon, H., Lebreton, J.-D., Metcalf, C.J.E., Neel, M.M., Parker, I.M., Takada, T., Valverde, T., Vélez-Espino, L.A., Wardle, G.M., Franco, M. & Vaupel, J.W. (2015) The COMPADRE Plant Matrix Database: An open online repository for plant demography. Journal of Ecology, 103, 202–218.

Salguero-Gómez, R., Jones, O.R., Jongejans, E., Blomberg, S.P., Hodgson, D.J., Mbeau-Ache, C., Zuidema, P.A., De Kroon, H. & Buckley, Y.M. (2016b) Fast–slow continuum and reproductive strategies structure plant life-history variation worldwide. Proceedings of the National Academy of Sciences, 113, 230–235.

Stearns, S.C. (1992) The Evolution of Life Histories, Oxford University Press, New York.

Steele, J.H., Brink, K.H. & Scott, B.E. (2019) Comparison of marine and terrestrial ecosystems: suggestions of an evolutionary perspective influenced by environmental variation. ICES Journal of Marine Science, 76, 50–59.

Strathmann, R.R. (1990) Why life histories evolve differently in the sea. American Zoologist, 30, 197–207.

Uhlenbeck, G.E. & Ornstein, L.S. (1930) On the Theory of the Brownian Motion. Phys. Rev., 36, 823–841.

Vermeij, G.J. & Grosberg, R.K. (2017) Rarity and persistence. Ecology Letters, 21, 3–8.

Webb, T.J. (2012) Marine and terrestrial ecology: Unifying concepts, revealing differences. Trends in Ecology and Evolution, 27, 535–541.

Webb, T.J. & Mindel, B.L. (2015) Global patterns of extinction risk in marine and non-marine systems. Current Biology, 25, 506–511.

Welch, B.L. (1947) The generalization of “Student’s” problem when several different population variances are involved. Biometrika, 34, 28–35.

Westoby, M. (1998) A leaf-height-seed (LHS) plant ecology strategy scheme. Plant and soil, 199, 213–227.

Winemiller, K.O. & Rose, K.A. (1992) Patterns of life-history diversification in North American fishes: implications for population regulation. Canadian Journal of Fisheries and aquatic sciences, 49, 2196–2218.

