## Appendix for "Aquatic and terrestrial organisms display contrasting life history strategies as a result of environmental adaptations"

### ****Appendix S2: Extended Methods****

*Data selection*

We selected only species in COMPADRE and COMADRE for which the MPMs: (i) were parameterised with field data from non-disturbed, unmanipulated populations to examine only natural populations; (ii) had at least three or more life cycle stages, as lower dimension MPMs typically lack the necessary quality for the estimation of life history traits (below; Salguero-Gómez & Plotkin, 2010); (iii) species for which the phylogeny is resolved, in order to be able to perform phylogenetic comparative analyses (Revell, 2010, 2012).

When MPMs existed for multiple populations within a given species and study, we calculated the arithmetic element-by-element mean MPM to obtain a single MPM per species (Tuljapurkar & Haridas, 2006; Salguero-Gómez *et al.*, 2016; Paniw *et al.*, 2018). When multiple studies existed for the same species, only studies with the longest study period were selected. This step ensured a better representation of the temporal variation that the natural populations experience.

These criteria filtered 684 terrestrial species and 117 aquatic species used in this study (Table S1). The number of species studied here represented a similar taxonomic coverage relative to the known biodiversity of the marine (0.05%), freshwater (0.02%) and terrestrial realm (0.04%; Table S1).

**Table S1. Comparison of the number of species examined in this study, compared to the number of currently accepted species worldwide, by taxonomic group and realm.** To contextualise the representativity of the examined species in this study across the tree of life, we compared the number of selected species with the currently known number of accepted species. We estimated the total number of species on land, freshwater and marine environments based on the rough estimates provided in Grosberg *et al.* (2012). We estimated the total number of species (*i.e.,* fraction of biodiversity) that our marine, freshwater and terrestrial species represented in comparison to the current species knowledge.

| Taxon | Studied marine species | Accepted marine species | Marine representativity (%) | Studied freshwater species | Accepted freshwater species | Freshwater representativity (%) | Studied terrestrial species | Accepted terrestrial species | Terrestrial representativity  (%) |
| --- | --- | --- | --- | --- | --- | --- | --- | --- | --- |
| Annelida | 0 | 9500 | 0.00 | 0 | 0 | 0 | 1 | 100 | 1.00 |
| Arthropoda | 2 | 9400 | 0.02 | 4 | 117000 | 0.00 | 4 | 6004500 | 0.00 |
| Chordata | 30 | 18625 | 0.16 | 35 | 22100 | 0.14 | 121 | 25000 | 0.51 |
| Cnidaria | 17 | 10000 | 0.17 | 0 | 50 | 0.00 | 0 | 0 | 0.00 |
| Echinodermata | 2 | 6500 | 0.03 | 0 | 0 | 0.00 | 0 | 0 | 0.00 |
| Mollusca | 11 | 61800 | 0.02 | 0 | 5900 | 0.00 | 0 | 27500 | 0.00 |
| Porifera | 3 | 8600 | 0.03 | 0 | 225 | 0.00 | 0 | 0 | 0.00 |
| Vascular macrophytes | 0 | 500 | 0.00 | 0 | 0 | 0.00 | 506 | 250000 | 0.20 |
| Non-vascular macrophytes | 13 | 10000 | 0.13 | 0 | 2500 | 0.00 | 1 | 30000 | 0.00 |
| **Total** | **78** | **171925** | **0.05** | **39** | **153875** | **0.02** | **638** | **8337500** | **0.01** |

*Life history traits*

For every species, we decomposed the MPM ***A*** into two sub-components (equation 1): the ***U*** matrix, which represents the survival-dependent vital rates (e.g. development, shrinkage, fission, etc); and the ***F*** matrix, containing the stage-specific per-capita reproduction rates (Caswell, 2001; Morris & Doak, 2002). Those species showing clonality were removed from the analyses, in order to avoid potential over estimation of survival rates. This decomposition facilitates the estimation of key life history traits such as the time elapsed since or to a given demographic event (e.g.*,* age at maturity, mean life expectancy; see Table S2).

***A*** = ***U*** + ***F***	equation 1

The traits *T*, *L_α_* and *R_0_* were calculated using stage-from-age demographic decompositions (Caswell 2001; p. 124-127; see Table S2), where the beginning of life was *a priori* defined as the first non-propagule stage in the life cycle of the organism (Burns *et al.*, 2010). This approach avoids uncertainties associated with the longevity of spores and seeds (Silvertown & Franco, 1993; Caswell, 2001; Burns *et al.*, 2010; Salguero-Gómez *et al.*, 2016) and assures the comparability with species without such life cycle stages. To calculate *S* and *H* (Demetrius, 1974; Keyfitz, 1977), we first obtained the age-specific survivorship curve (*l_x_*), and the age-specific fertility trajectory (*m_x_*) following Caswell (2001; p. 118-121), and implemented the formulae described in Table S2. The traits progressive development (*γ*) and sexual reproduction (*φ*) summarise investments on development and reproduction annually for all stages across the life cycle weighted by the relative representation of stages under stationary conditions (Table S2).

**Table S2. Formulation of the life history traits used to explore the variation in life history strategies in the studied 638 terrestrial and 117 aquatic species.** *λ* is the deterministic population growth rate, which corresponds to the dominant eigenvalue of the matrix ***A*** (Caswell, 2001); *l_x_* and *m_x_* are the age-specific survival and fertility schedules, respectively; ***U*** and ***F*** are the sub-matrices of survival- and fertility-dependent processes (equation 1); ***U’*** is the survival-independent matrix of transition probabilities (equation 2); ***w*** is the stable stage distribution of the matrix ***A***, and *i* and *j* are the row and column entries of the matrix population model, respectively.

|  | **Life history trait** | | **Definition** | **Calculation** |
| --- | --- | --- | --- | --- |
| ***Turnover*** | Generation time | *T* | Number of years required for an average individual in the population to replace itself. | $T=\frac{\sum x\times\left( l_{x}\times m_{x} \right)}{\sum\left( l_{x}\times m_{x} \right)}$ |
| ***Survival*** | Rate of senescence | *H* | Shape of the age-specific survivorship curve *l_x_* as quantified by Keyfitz’ entropy (*H*).  H >1, = 1, <1 correspond to species whose mortality hazards decrease, stay constant, or increase with age, respectively. | $H=\frac{-\log(l_{x})l_{x}}{\sum l_{x}}$ |
|  | Age at maturity | *L_α_* | Average amount of time from birth to reproduction. | Caswell 2001, p. 124 |
| ***Development*** | Mean vital rate of progressive growth (γ) | *γ* | Mean probability of transitioning forward to a larger/more developed stage in the life cycle of the species, weighted by the stable stage distribution, ***w***. | $\gamma=\sum_{1}^{m} \left. \bar{U'}_{i,j}\bar{w}_{j} \right\vert_{i<j}$ |
| ***Reproduction*** | Mean vital rate of sexual reproduction | *φ* | Mean per-capita number of sexual recruits across stages in the life cycle of the species, weighted by ***w***. | $\phi=\sum_{1}^{m} \bar{F}_{j}\bar{w}_{j}$ |
|  | Degree of iteroparity | *S* | Temporal spread of reproduction throughout lifespan as quantified by Demetrius’ (1974) entropy (*S*). High/low *S* values correspond to iteroparous/semelparous populations. | $S=-e^{-log\lambda}l_{x}m_{x}log\left( e^{-log\lambda}l_{x}m_{x} \right)$ |

*Missing data*

Some of the life history traits were not possible to calculate with the available demographic data in COMPADRE and COMADRE. We explored the proportion of missing values for species in both terrestrial and the aquatic realms (Figure S1). In the aquatic environment, for most of the traits, the proportion of missing values was higher than for the terrestrial ones (Figure S1). However, for both datasets the trait with the highest proportion of missing values was the rate of senescence (*H*), with values below 45% for aquatic species and below 30% in terrestrial species (Figure S1).

The proportion of missing values changed according to the taxonomic grups (Table S3). Despite the higher proportion of missing values in Aquatic species (Figure S1), the taxonomic groups showing a higher proportion of missing values were mostly terrestrial (Table S3). The terrestrial species of the Class Bryopsida, Clitellata and Pteridopsida, and the aquatic species of the Class Thaliacea, showed a 100% of missing values for more than two traits (Table S3). Corals (Anthozoa), amphibians and insects also showed a high proportion of missing values for the rate of senescence (Table S3).

Because the pPCA analyses require complete datasets, we imputed the missing data using function amelia from the Amelia package (Honaker *et al.*, 2011), as detailed in the Methods’ section *Exploring dominant axes of life history strategies* of the main manuscript.

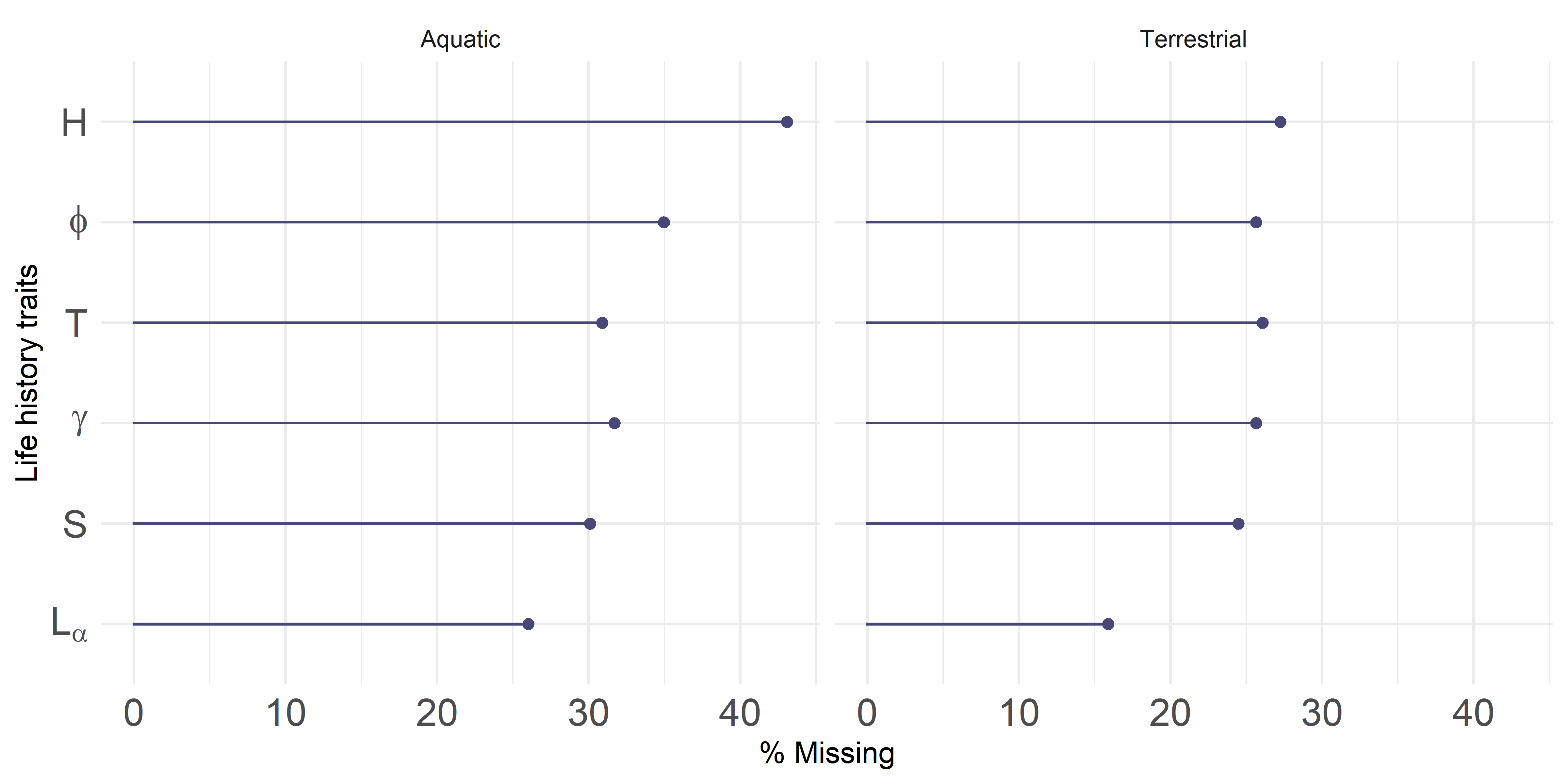

**Figure S1. Proportion of missing life history trait values for the species included in our analyses per realm.** The missing values were estimated from the overall data set. The symbols represent, in descendent order, rate of senescence (H), mean vital rate of sexual reproduction (φ), generation time (T), mean vital rate of growth rate (γ), degree of iteroparity (S) and age at maturity (L_α_).

**Table S3. Proportion of missing values for each of the life history traits of the life history traits used to explore the variation in life history strategies in the studied 638 terrestrial and 117 aquatic species.** classified according to the realm and taxonomic Class.

| **Realm** | **Class** | **Generation time** | **Rate of senescence** | **Age at sexual maturity** | **Mean vital rate of growth** | **Mean vital rate of reproduction** | **Degree of iteroparity** |
| --- | --- | --- | --- | --- | --- | --- | --- |
| Aquatic | Actinopterygii | 11.76 | 32.35 | 11.76 | 11.76 | 11.76 | 11.76 |
| Aquatic | Amphibia | 0.00 | 100.00 | 0.00 | 0.00 | 0.00 | 0.00 |
| Aquatic | Anthozoa | 29.41 | 82.35 | 29.41 | 29.41 | 29.41 | 29.41 |
| Aquatic | Aves | 0.00 | 0.00 | 0.00 | 0.00 | 0.00 | 0.00 |
| Aquatic | Bivalvia | 33.33 | 33.33 | 16.67 | 33.33 | 33.33 | 33.33 |
| Aquatic | Branchiopoda | 0.00 | 100.00 | 0.00 | 0.00 | 0.00 | 0.00 |
| Aquatic | Demospongiae | 0.00 | 0.00 | 0.00 | 0.00 | 0.00 | 0.00 |
| Aquatic | Echinoidea | 0.00 | 100.00 | 0.00 | 0.00 | 0.00 | 0.00 |
| Aquatic | Elasmobranchii | 0.00 | 0.00 | 0.00 | 0.00 | 0.00 | 0.00 |
| Aquatic | Florideophyceae | 66.67 | 33.33 | 33.33 | 33.33 | 33.33 | 33.33 |
| Aquatic | Gastropoda | 20.00 | 20.00 | 20.00 | 20.00 | 20.00 | 20.00 |
| Aquatic | Insecta | 0.00 | 0.00 | 0.00 | 0.00 | 0.00 | 0.00 |
| Aquatic | Malacostraca | 0.00 | 0.00 | 0.00 | 0.00 | 0.00 | 0.00 |
| Aquatic | Mammalia | 30.77 | 53.85 | 23.08 | 30.77 | 30.77 | 30.77 |
| Aquatic | Maxillopoda | 0.00 | 0.00 | 0.00 | 0.00 | 0.00 | 0.00 |
| Aquatic | Phaeophyceae | 30.00 | 20.00 | 10.00 | 30.00 | 30.00 | 20.00 |
| Aquatic | Reptilia | 12.50 | 37.50 | 0.00 | 12.50 | 12.50 | 12.50 |
| Aquatic | Thaliacea | 100.00 | 100.00 | 0.00 | 0.00 | 0.00 | 100.00 |
| Terrestrial | Asterids | 0.00 | 0.00 | 0.00 | 0.00 | 0.00 | 0.00 |
| Terrestrial | Aves | 11.63 | 13.95 | 6.98 | 13.95 | 13.95 | 11.63 |
| Terrestrial | Bryopsida | 100.00 | 100.00 | 0.00 | 100.00 | 0.00 | 100.00 |
| Terrestrial | Clitellata | 100.00 | 100.00 | 0.00 | 100.00 | 0.00 | 100.00 |
| Terrestrial | Cycadopsida | 25.00 | 25.00 | 12.50 | 25.00 | 25.00 | 25.00 |
| Terrestrial | Insecta | 25.00 | 75.00 | 25.00 | 25.00 | 25.00 | 25.00 |
| Terrestrial | Liliopsida | 30.08 | 30.08 | 25.56 | 30.08 | 30.08 | 30.08 |
| Terrestrial | Magnoliopsida | 25.52 | 24.33 | 12.17 | 24.33 | 24.33 | 23.15 |
| Terrestrial | Mammalia | 25.76 | 42.42 | 21.21 | 25.76 | 25.76 | 25.76 |
| Terrestrial | Pinopsida | 29.63 | 29.63 | 18.52 | 29.63 | 29.63 | 29.63 |
| Terrestrial | Polypodiopsida | 0.00 | 0.00 | 0.00 | 0.00 | 0.00 | 0.00 |
| Terrestrial | Pteridopsida | 100.00 | 100.00 | 0.00 | 0.00 | 0.00 | 100.00 |
| Terrestrial | Reptilia | 11.76 | 17.65 | 11.76 | 11.76 | 11.76 | 11.76 |

### Appendix S3: Extended phylogenetic methods

To build the phylogenetic tree with the species included in our analyses we used data from Open Tree of Life (OTL, https://tree.opentreeoflife.org; Hinchliff *et al.*, 2015). OTL is an assembly of published phylogenies, together with taxonomic classifications (Hinchliff *et al.*, 2015), enabling us to account for the phylogenetic relatedness among the species included in our study. To build the tree we first checked that the species names in our data were taxonomically accepted using the *taxize* R package (Chamberlain & Szöcs, 2013)*.* Then we obtained algae, plant, and animal phylogenetic trees from OTL for the list of species in our data using the *rotl* R package (Michonneau *et al.*, 2016). Finally, we created a supertree using the function *bind.tree* in the *phytools* package (Revell, 2012), assembling first plants and algae trees, and then binding the animal tree with sponges (X*estospongia muta*, giant barrel sponge) as the outgroup.

Because OTL combines taxonomic and phylogenetic information to account for the phylogenetic relatedness of species we computed the branch lengths and resolved polytomies (Revell, 2010, 2012). The branch length of the resulting tree was computed using the *compute.brlen* function from the R package *ape* (Paradis *et al.*, 2004), with Grafen’s arbitrary branch lengths (Grafen, 1989). Polytomies (*i.e.*, a node in the tree with >2 species with a common immediate ancestor) were resolved using the function *multi2di* from the *ape* package (Paradis *et al.*, 2004). Briefly, this function transforms polytomies into a series of random dichotomies with one or several branches of length zero. To determine the sensitivity of our results to the choice of a particular set of arbitrary branch lengths, we repeated the our analyses setting all the branch lengths to one and using Pagel’s branch length using the softward Mesquite 1.05 (Maddison & Maddison, 2018) and its PDAP module 1.06 (Midford *et al.*, 2005). We did not observe substantial changes in our findings (see Tables S4-S7).

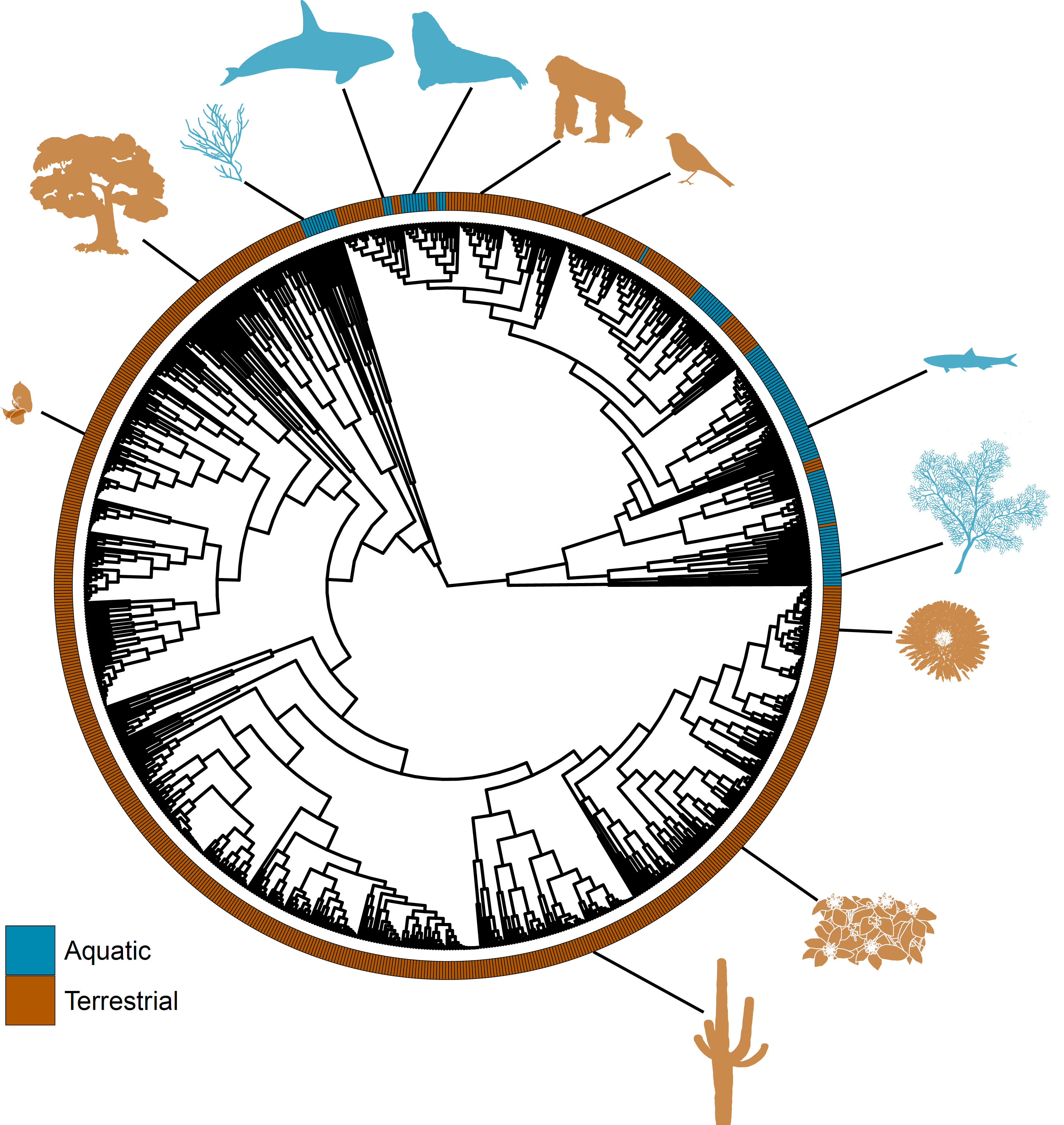

**Figure S2. Phylogeny included in our comparative analyses.** Species-level phylogenetic tree with data from Open Tree of Life (OTL, <https://tree.opentreeoflife.org> (Hinchliff *et al.*, 2015), for the 117 aquatic (blue) and 638 terrestrial (brown) species included in our analyses. The silhouettes, starting at the bottom right, and counter-clock-wise, correspond to: *Pseudomitrocereus fulviceps,* *Lantana camara*, *Solidago mollis*, *Paramuricea clavata*, *Clinostomus funduloides*, *Setophaga cerulea*, *Gorilla beringei*, *Mirounga leonina*, *Orcinus orca*, *Gracilaria gracilis*, *Chlorocardium rodiei* and *Cypripedium calceolus*.

**Table S4. Loadings of phylogenetically corrected principal component analysis (pPCA) equalizing the evolutionary relationships between species by setting the branch lengths of the phylogeny to one** (see Methods in the main manuscript)**.** Headers of axes with associated eigenvalues above 1, indicating retention to explain observed variation (Legendre & Legendre, 2012), are in bold. Bold numbers indicate loading absolute values >0.50. Here are presented the mean values and the standard errors of the values coming from the multiple imputed data sets. See Table S2 for more details about the life history traits.

| **Life history traits** | | **PC1** | **PC2** | PC3 | PC4 | PC5 | PC6 |
| --- | --- | --- | --- | --- | --- | --- | --- |
| **Turnover** | Generation time (*T*) | **0.83±0.00** | -0.08±0.01 | -0.23±0.01 | -0.32±0.01 | 0.19±0.01 | 0.33±0.00 |
| **Survival** | Rate of senescence (*H*) | **0.72±0.01** | 0.24±0.01 | -0.24±0.01 | 0.59±0.01 | -0.07±0.01 | 0.08±0.00 |
|  | Age at maturity (*L_α_*) | **0.80±0.00** | 0.01±0.01 | -0.43±0.01 | -0.23±0.01 | -0.16±0.01 | -0.32±0.00 |
| **Development** | Development (*γ*) | **-0.73±0.00** | -0.11±0.01 | -0.54±0.01 | 0.11±0.01 | 0.37±0.01 | -0.06±0.00 |
| **Reproduction** | Mean reproducitve output (*Φ*) | **-0.69±0.01** | **0.51±0.01** | -0.34±0.01 | -0.13±0.01 | -0.33±0.01 | 0.15±0.00 |
|  | Degree of iteroparity (*S*) | 0.18±0.02 | **0.92±0.00** | 0.17±0.01 | -0.08±0.01 | 0.26±0.01 | -0.09±0.00 |
| **Eigenvalue** | | **3.08±0.03** | **1.36±0.01** | 0.78±0.01 | 0.58±0.01 | 0.42±0.01 | 0.27±0.01 |
| **Proportion of variance explained** | | 47.42±0.34% | 21.02±0.11% | 12.05±0.17% | 8.94±0.12% | 6.45±0.09% | 4.11±0.05% |
| **Cumulative proportion of variance explained** | | 47.42% | 68.44% | 80.50% | 89.44% | 95.89% | 100% |
| **Pagel’s λ** | | 0.26±0.01 | | | | | |

**Table S5. Phylogenetically Generalised Least Squares (PGLS) regressions between life history traits of aquatic and terrestrial taxa with imputed data assuming a Brownian motion evolutionary model, equalizing the evolutionary relationships between species by setting the branch lengths of the phylogeny to one** (see Methods in the main manuscript). Outputs correspond to the mean values of 10 imputed data sets: slopes, the effect of the realm, and the interaction of the realm with the second trait. *P*-adjusted is the adjusted *P*-value after a Bonferroni correction.

| Traits | Slope | SD | *t*-value | *P* | *P*-adjusted | Realm | SD | *t*-value | *P* | *P*-adjusted | Pagel’s *λ* | CI | R^2^ |
| --- | --- | --- | --- | --- | --- | --- | --- | --- | --- | --- | --- | --- | --- |
| ***T ~ H*** | **1.59** | **0.19** | **8.26** | **<0.001** | **<0.001** | 0.08 | 0.38 | 0.20 | 0.83 | 1.00 | 0.51 | 0.95 | 0.14 |
| ***T ~ Lα*** | **0.88** | **0.05** | **16.45** | **<0.001** | **<0.001** | 0.05 | 0.20 | 0.25 | 0.78 | 1.00 | 0.05 | 0.95 | 0.38 |
| ***T ~ γ*** | **-4.31** | **0.55** | **-7.81** | **<0.001** | **<0.001** | 0.14 | 0.42 | 0.34 | 0.74 | 1.00 | 0.64 | 0.95 | 0.12 |
| ***T ~ φ*** | **-6.42** | **0.64** | **-9.99** | **<0.001** | **<0.001** | -0.01 | 0.37 | -0.02 | 0.89 | 1.00 | 0.54 | 0.95 | 0.19 |
| *T ~ S* | 0.06 | 0.09 | 0.67 | 0.50 | 0.56 | 0.29 | 0.48 | 0.62 | 0.54 | 1.00 | 0.72 | 0.95 | <0.001 |
| ***H ~ Lα*** | **0.15** | **0.01** | **10.41** | **<0.001** | **<0.001** | 0.06 | 0.07 | 0.89 | 0.38 | 1.00 | 0.33 | 0.95 | 0.20 |
| ***H ~ γ*** | **-1.22** | **0.15** | **-8.26** | **<0.001** | **<0.001** | 0.06 | 0.12 | 0.53 | 0.60 | 1.00 | 0.66 | 0.95 | 0.14 |
| ***H ~ φ*** | **-1.04** | **0.19** | **-5.38** | **<0.001** | **<0.001** | 0.05 | 0.13 | 0.48 | 0.63 | 1.00 | 0.69 | 0.95 | 0.06 |
| *H ~ S* | 0.07 | 0.02 | 3.39 | 0.07 | 0.07 | 0.08 | 0.14 | 0.68 | 0.50 | 1.00 | 0.70 | 0.95 | 0.03 |
| ***Lα ~ γ*** | **-2.44** | **0.33** | **-5.77** | **<0.001** | **<0.001** | 0.17 | 0.34 | 0.33 | 0.56 | 1.00 | 0.83 | 0.95 | 0.13 |
| ***Lα ~ φ*** | **-2.61** | **0.40** | **-6.25** | **<0.001** | **<0.001** | 0.05 | 0.34 | 0.03 | 0.81 | 1.00 | 0.84 | 0.95 | 0.08 |
| *Lα ~ S* | -0.02 | 0.05 | -0.33 | 0.67 | 0.69 | 0.15 | 0.39 | 0.25 | 0.64 | 1.00 | 0.89 | 0.95 | <0.001 |
| ***γ ~ φ*** | **0.49** | **0.05** | **10.13** | **<0.001** | **<0.001** | -0.03 | 0.03 | -1.33 | 0.18 | 1.00 | 0.34 | 0.95 | 0.20 |
| ***γ ~ S*** | **<0.001** | **0.01** | **-2.87** | **<0.001** | **<0.001** | -0.03 | 0.04 | -1.00 | 0.31 | 1.00 | 0.44 | 0.95 | 0.05 |
| ***φ ~ S*** | **0.04** | **0.01** | **6.08** | **<0.001** | **<0.001** | -0.04 | 0.03 | -1.36 | 0.17 | 1.00 | 0.59 | 0.95 | 0.08 |

**Table S6. Loadings of phylogenetically corrected principal component analysis (pPCA) using Pagel’s branch length** (see Methods in the main manuscript). Headers of axes with associated eigenvalues >1, indicating retention to explain observed variation (Legendre & Legendre, 2012), are shown in bold. Bold numbers indicate loading absolute values >0.50. The mean values and associated standard errors were obtained from 10 imputed data sets. See Table S2 for more details about the life history traits.

| **Life history traits** | | **PC1** | **PC2** | PC3 | PC4 | PC5 | PC6 |
| --- | --- | --- | --- | --- | --- | --- | --- |
| **Turnover** | Generation time (*T*) | **0.74±0.01** | -0.03±0.02 | -0.39±0.03 | 0.26±0.09 | 0.12±0.03 | 0.34±0.01 |
| **Survival** | Rate of senescence (*H*) | **0.64±0.01** | 0.41±0.01 | -0.27±0.04 | -0.46±0.10 | 0.04±0.04 | 0.06±0.01 |
|  | Age at maturity (*L_α_*) | **0.73±0.01** | 0.07±0.02 | -0.41±0.02 | 0.16±0.07 | -0.25±0.02 | -0.40±0.01 |
| **Development** | Development (*γ*) | **-0.74±0.01** | -0.21±0.02 | **-0.53±0.02** | -0.06±0.03 | 0.31±0.02 | -0.09±0.01 |
| **Reproduction** | Mean reproducitve output (*Φ*) | **-0.74±0.01** | **0.54±0.01** | -0.22±0.01 | 0.04±0.04 | -0.30±0.01 | 0.11±0.00 |
|  | Degree of iteroparity (*S*) | 0.06**±0.02** | **0.86±0.01** | 0.19±0.01 | 0.19±0.05 | 0.36±0.02 | -0.11±0.01 |
| **Eigenvalue** | | **2.97±0.17** | **1.48±0.13** | 0.91±0.11 | 0.65±0.05 | 0.50±0.01 | 0.30±0.01 |
| **Proportion of variance explained** | | 43.96±0.84% | 21.58±0.29% | 13.22±0.43% | 9.53±0.17% | 7.30±0.20% | 4.40±0.10% |
| **Cumulative proportion of variance explained** | | 43.96% | 65.54% | 78.77% | 88.30% | 95.60% | 100% |
| **Pagel’s λ** | | 0.93±0.01 | | | | | |

**Table S7. Phylogenetically Generalised Least Squares (PGLS) regressions between life history attributes for aquatic and terrestrial taxa with 10 imputed data sets, assuming a Brownian motion evolutionary using Pagel’s branch length** (see Methods in the main manuscript). Outputs correspond to the mean values of 10 imputed data sets: slopes, the effect of the realm, and the interaction of the realm with the second trait. *P*-adjusted is the adjusted *P*-value after a Bonferroni correction.

| Traits | Slope | SD | *t*-value | *P* | *P*-adjusted | Realm | SD | *t*-value | *P* | *P*-adjusted | Pagel’s *λ* | CI | R^2^ |
| --- | --- | --- | --- | --- | --- | --- | --- | --- | --- | --- | --- | --- | --- |
| ***T ~ H*** | **0.99** | **0.18** | **5.38** | **<0.001** | **<0.001** | 0.45 | 0.27 | 1.63 | 0.11 | 1.00 | 0.92 | 0.95 | 0.07 |
| ***T ~ Lα*** | **0.78** | **0.04** | **19.27** | **<0.001** | **<0.001** | 0.25 | 0.15 | 1.70 | 0.09 | 1.00 | 0.51 | 0.95 | 0.46 |
| ***T ~ γ*** | **-2.89** | **0.39** | **-7.53** | **<0.001** | **<0.001** | 0.52 | 0.25 | 2.04 | 0.04 | 1.00 | 0.94 | 0.95 | 0.12 |
| ***T ~ φ*** | **-4.64** | **0.44** | **-10.03** | **<0.001** | **<0.001** | 0.32 | 0.24 | 1.35 | 0.18 | 1.00 | 0.93 | 0.95 | 0.20 |
| *T ~ S* | -0.06 | 0.09 | -0.25 | 0.39 | 0.44 | 0.57 | 0.28 | 2.02 | 0.05 | 1.00 | 0.95 | 0.95 | 0.01 |
| ***H ~ Lα*** | **-0.44** | **0.08** | **2.55** | **<0.001** | **<0.001** | 0.15 | 0.10 | 1.63 | 0.14 | 0.98 | 0.93 | 0.95 | 0.11 |
| ***H ~ γ*** | **-1.23** | **0.17** | **-7.57** | **<0.001** | **<0.001** | 0.16 | 0.12 | 1.56 | 0.16 | 1.00 | 0.97 | 0.95 | 0.12 |
| ***H ~ φ*** | **-0.77** | **0.16** | **-3.50** | **0.02** | **0.02** | 0.14 | 0.13 | 1.30 | 0.23 | 1.00 | 0.97 | 0.95 | 0.05 |
| *H ~ S* | 0.06 | 0.03 | 2.90 | 0.07 | 0.08 | 0.18 | 0.13 | 1.50 | 0.16 | 1.00 | 0.97 | 0.95 | 0.04 |
| ***Lα ~ γ*** | **-1.13** | **0.19** | **-1.01** | **<0.001** | **<0.001** | 0.14 | 0.15 | 0.80 | 0.44 | 1.00 | 0.97 | 0.95 | 0.15 |
| *Lα ~ φ* | -0.85 | 0.15 | -2.38 | 0.11 | 0.12 | 0.12 | 0.16 | 0.55 | 0.58 | 1.00 | 0.98 | 0.95 | 0.07 |
| *Lα ~ S* | -0.02 | 0.03 | -0.15 | 0.26 | 0.29 | 0.17 | 0.17 | 0.46 | 0.43 | 1.00 | 0.99 | 0.95 | 0.01 |
| ***γ ~ φ*** | **0.27** | **0.03** | **6.86** | **<0.001** | **<0.001** | 0.01 | 0.04 | -0.02 | 0.40 | 0.93 | 0.89 | 0.95 | 0.15 |
| *γ ~ S* | 0.02 | 0.01 | 0.34 | 0.12 | 0.12 | 0.03 | 0.05 | 0.29 | 0.68 | 0.87 | 0.83 | 0.95 | 0.05 |
| ***φ ~ S*** | **0.04** | **0.01** | **5.02** | **<0.001** | **<0.001** | -0.06 | 0.03 | -1.89 | 0.06 | 1.00 | 0.99 | 0.95 | 0.06 |

### Appendix S4: Extended results

**Table S8. Loadings of phylogenetically corrected principal component analysis (pPCA) by the life history traits, grouped into turnover, survival, development and reproduction attributes for the 117 aquatic and 638 terrestrial species in this study** **(see Table S2)**. Headers of axes with associated eigenvalues >1, indicating retention to explain observed variation (Legendre & Legendre, 2012), are shown in bold. Bold numbers indicate loading absolute values >0.50. Outputs correspond to the mean values of 10 imputed data sets.

| **Life history traits** | | **PC1** | **PC2** | PC3 | PC4 | PC5 | PC6 |
| --- | --- | --- | --- | --- | --- | --- | --- |
| **Turnover** | Generation time (*T*) | 0.83±0.00 | -0.08±0.01 | -0.23±0.01 | -0.32±0.01 | 0.19±0.01 | 0.33±0.00 |
| **Survival** | Rate of senescence (*H*) | 0.72±0.01 | 0.24±0.01 | -0.24±0.01 | 0.59±0.01 | -0.07±0.01 | 0.08±0.00 |
|  | Age at maturity (*L_α_*) | 0.80±0.00 | 0.01±0.01 | -0.43±0.01 | -0.23±0.01 | -0.16±0.01 | -0.32±0.00 |
| **Development** | Development (*γ*) | -0.73±0.00 | -0.11±0.01 | -0.54±0.01 | 0.11±0.01 | 0.37±0.01 | -0.06±0.00 |
| **Reproduction** | Mean reproducitve output (*Φ*) | -0.69±0.01 | 0.51±0.01 | -0.34±0.01 | -0.13±0.01 | -0.33±0.01 | 0.15±0.00 |
|  | Degree of iteroparity (*S*) | 0.18±0.02 | 0.92±0.00 | 0.17±0.01 | -0.08±0.01 | 0.26±0.01 | -0.09±0.00 |
| **Eigenvalue** | | **3.08±0.03** | **1.36±0.01** | 0.78±0.01 | 0.58±0.01 | 0.42±0.01 | 0.27±0.01 |
| **Proportion of variance explained** | | 47.42±0.34% | 21.02±0.11% | 12.05±0.17% | 8.94±0.12% | 6.45±0.09% | 4.11±0.05% |
| **Cumulative proportion of variance explained** | | 47.42% | 68.44% | 80.50% | 89.44% | 95.89% | 100% |
| **Pagel’s λ** | | 0.26±0.01 | | | | | |

**Table S9. Phylogenetically Generalised Least Squares (PGLS) regressions between life history attributes for aquatic and terrestrial taxa with imputed data (n=638 terrestrial, n=117 aquatic species)**. Outputs correspond to the mean values of 10 imputed data sets: slopes, the effect of the realm and the interaction of the realm with the second trait. *P*-adjusted is the adjusted *P*-value after a Bonferroni correction (see Table S2).

| Traits | Slope | SD | *t*-value | *P* | *P*-adjusted | Realm | SD | *t*-value | *P* | *P*-adjusted | Pagel’s *λ* | CI | R^2^ |
| --- | --- | --- | --- | --- | --- | --- | --- | --- | --- | --- | --- | --- | --- |
| *T ~ H* | **1.81** | **0.17** | **10.50** | **<0.001** | **<0.001** | 0.12 | 0.25 | 0.49 | 0.63 | 1.00 | 0.45 | 0.95 | 0.18 |
| *T ~ Lα* | **0.91** | **0.05** | **18.36** | **<0.001** | **<0.001** | 0.16 | 0.17 | 0.95 | 0.35 | 1.00 | 0.11 | 0.95 | 0.40 |
| *T ~ γ* | **-4.09** | **0.51** | **-8.06** | **<0.001** | **<0.001** | 0.22 | 0.28 | 0.77 | 0.44 | 1.00 | 0.61 | 0.95 | 0.12 |
| *T ~ φ* | **-5.81** | **0.53** | **-10.93** | **<0.001** | **<0.001** | 0.06 | 0.27 | 0.23 | 0.82 | 1.00 | 0.61 | 0.95 | 0.20 |
| *T ~ S* | 0.01 | 0.08 | 0.13 | 0.89 | 0.91 | 0.44 | 0.32 | 1.40 | 0.16 | 1.00 | 0.71 | 0.95 | <0.001 |
| *H ~ Lα* | **0.16** | **0.01** | **11.83** | **<0.001** | **<0.001** | 0.11 | 0.06 | 2.02 | 0.05 | 0.86 | 0.42 | 0.95 | 0.23 |
| *H ~ γ* | **-0.99** | **0.12** | **-8.31** | **<0.001** | **<0.001** | 0.10 | 0.07 | 1.42 | 0.17 | 1.00 | 0.65 | 0.95 | 0.13 |
| *H ~ φ* | **-0.70** | **0.14** | **-5.18** | **<0.001** | **<0.001** | 0.10 | 0.07 | 1.38 | 0.18 | 1.00 | 0.68 | 0.95 | 0.06 |
| *H ~ S* | **0.08** | **0.02** | **4.47** | **<0.001** | **<0.001** | 0.13 | 0.07 | 1.78 | 0.09 | 0.96 | 0.74 | 0.95 | 0.05 |
| *Lα ~ γ* | **-2.47** | **0.35** | **-7.08** | **<0.001** | **<0.001** | 0.09 | 0.22 | 0.40 | 0.69 | 1.00 | 0.82 | 0.95 | 0.09 |
| *Lα ~ φ* | **-2.69** | **0.38** | **-7.01** | **<0.001** | **<0.001** | <0.001 | 0.23 | 0.02 | 0.97 | 1.00 | 0.83 | 0.95 | 0.09 |
| *Lα ~ S* | -0.02 | 0.05 | -0.45 | 0.65 | 0.73 | 0.21 | 0.25 | 0.85 | 0.40 | 1.00 | 0.87 | 0.95 | <0.001 |
| *γ ~ φ* | **0.47** | **0.04** | **10.50** | **<0.001** | **<0.001** | -0.03 | 0.02 | -1.25 | 0.21 | 1.00 | 0.46 | 0.95 | 0.19 |
| *γ ~ S* | **-0.02** | **0.01** | **-3.18** | **<0.001** | **<0.001** | -0.04 | 0.03 | -1.73 | 0.08 | 1.00 | 0.64 | 0.95 | 0.03 |

**Table S10. Loadings of phylogenetically corrected principal component analysis (pPCA) without excluding outliers (see Table S2).** Headers of axes with associated eigenvalues >1, indicating retention to explain observed variation (Legendre & Legendre 2012), are shown in bold. Bold numbers indicate loading absolute values >0.50. Outputs correspond to the mean values of 10 imputed data sets.

| **Life history traits** | | **PC1** | **PC2** | **PC3** | PC4 | PC5 | PC6 |
| --- | --- | --- | --- | --- | --- | --- | --- |
| **Turnover** | Generation time (*T*) | **0.85±0.00** | -0.05±0.01 | -0.06±0.03 | -0.17±0.01 | -0.01±0.10 | 0.36±0.00 |
| **Survival** | Rate of senescence (*H*) | **0.73±0.00** | 0.20±0.01 | -0.06±0.03 | -0.05±0.03 | 0.00±0.21 | 0.06±0.01 |
|  | Age at maturity (*L_α_*) | **0.82±0.00** | 0.05±0.01 | -0.11±0.05 | -0.34±0.01 | 0.00±0.07 | -0.36±0.00 |
| **Development** | Development (*γ*) | **-0.40±0.01** | -0.13±0.03 | **-0.54±0.24** | -0.04±0.01 | 0.01±0.01 | 0.03±0.00 |
| **Reproduction** | Mean reproducitve output (*Φ*) | **-0.60±0.01** | **0.52±0.01** | 0.06±0.03 | -0.59±0.01 | -0.01±0.02 | 0.08±0.00 |
|  | Degree of iteroparity (*S*) | 0.14±0.01 | **0.89±0.01** | -0.09±0.05 | 0.37±0.01 | 0.00±0.05 | -0.02±0.00 |
| **Eigenvalue** | | **2.61±0.01** | **1.27±0.01** | **1.08±0.01** | 0.72±0.01 | 0.60±0.01 | 0.28±0.01 |
| **Proportion of variance explained** | | 39.81±0.27% | 19.42±0.08% | 16.50±0.10% | 10.91±0.13% | 9.08±0.12% | 4.28±0.04% |
| **Cumulative proportion of variance explained** | | 39.81% | 59.22% | 75.73% | 86.64% | 95.72% | 100% |
| **Pagel’s λ** | | 0.25±0.01 | | | | | |

**Table S11. Phylogenetically Generalised Least Squares (PGLS) regressions between life history attributes for aquatic and terrestrial taxa with imputed data and including outliers** Outputs correspond to the mean values of 10 imputed data sets: slopes, the effect of the realm and the interaction of the realm with the second trait. *P*-adjusted is the adjusted *P*-value after a Bonferroni correction.

| Traits | Slope | SD | *t*-value | *P* | *P*-adjusted | Realm | SD | *t*-value | *P* | *P*-adjusted | Pagel’s *λ* | CI | R^2^ |
| --- | --- | --- | --- | --- | --- | --- | --- | --- | --- | --- | --- | --- | --- |
| *T ~ H* | **1.48** | **0.18** | **8.02** | **<0.001** | **<0.001** | 0.20 | 0.41 | 0.49 | 0.63 | 1.00 | 0.55 | 0.95 | 0.12 |
| *T ~ Lα* | **0.82** | **0.05** | **17.12** | **<0.001** | **<0.001** | -0.02 | 0.21 | -0.09 | 0.87 | 1.00 | 0.06 | 0.95 | 0.39 |
| *T ~ γ* | **-3.69** | **0.45** | **-6.33** | **<0.001** | **<0.001** | 0.12 | 0.39 | 0.28 | 0.78 | 1.00 | 0.61 | 0.95 | 0.13 |
| *T ~ φ* | **-1.11** | **0.19** | **-6.09** | **<0.001** | **<0.001** | 0.21 | 0.43 | 0.46 | 0.65 | 1.00 | 0.67 | 0.95 | 0.07 |
| *T ~ S* | 0.06 | 0.09 | 0.44 | 0.36 | 0.42 | 0.24 | 0.45 | 0.50 | 0.62 | 1.00 | 0.70 | 0.95 | <0.001 |
| *T ~ R_0_* | <0.001 | 0.04 | 0.12 | 0.79 | 0.79 | 0.26 | 0.46 | 0.53 | 0.60 | 1.00 | 0.71 | 0.95 | <0.001 |
| *H ~ Lα* | **0.12** | **0.01** | **9.39** | **0.01** | **0.01** | <0.001 | 0.11 | <0.001 | 0.94 | 1.00 | 0.72 | 0.95 | 0.17 |
| *H ~ γ* | **-1.13** | **0.14** | **-8.08** | **<0.001** | **<0.001** | 0.03 | 0.14 | 0.11 | 0.90 | 1.00 | 0.72 | 0.95 | 0.12 |
| *H ~ φ* | **-0.17** | **0.05** | **-3.08** | **<0.001** | **<0.001** | 0.04 | 0.15 | 0.21 | 0.84 | 1.00 | 0.76 | 0.95 | 0.02 |
| *H ~ S* | 0.05 | 0.03 | 2.47 | 0.09 | 0.09 | 0.03 | 0.15 | 0.17 | 0.87 | 1.00 | 0.78 | 0.95 | 0.01 |
| *H ~ R_0_* | 0.03 | 0.01 | 1.99 | 0.06 | 0.07 | 0.04 | 0.15 | 0.25 | 0.80 | 1.00 | 0.77 | 0.95 | 0.01 |
| *Lα ~ γ* | **-2.43** | **0.33** | **-6.13** | **<0.001** | **<0.001** | 0.17 | 0.35 | 0.34 | 0.58 | 1.00 | 0.77 | 0.95 | 0.10 |
| *Lα ~ φ* | **-0.39** | **0.12** | **-3.18** | **<0.001** | **<0.001** | 0.16 | 0.38 | 0.30 | 0.64 | 1.00 | 0.81 | 0.95 | 0.02 |
| *Lα ~ S* | -0.01 | 0.06 | -0.23 | 0.82 | 0.84 | 0.17 | 0.40 | 0.30 | 0.62 | 1.00 | 0.82 | 0.95 | <0.001 |
| *Lα ~ R_0_* | **0.09** | **0.03** | **3.28** | **<0.001** | **<0.001** | 0.18 | 0.39 | 0.38 | 0.65 | 1.00 | 0.77 | 0.95 | 0.02 |
| *γ ~ φ* | **0.08** | **0.02** | **4.94** | **<0.001** | **<0.001** | -0.03 | 0.03 | -1.05 | 0.30 | 1.00 | 0.33 | 0.95 | 0.05 |
| *γ ~ S* | **-0.01** | **0.01** | **-2.25** | **<0.001** | **<0.001** | -0.01 | 0.04 | -0.70 | 0.35 | 1.00 | 0.37 | 0.95 | 0.03 |
| *γ ~ R_0_* | <0.001 | <0.001 | -0.68 | 0.49 | 0.54 | -0.03 | 0.03 | -1.02 | 0.31 | 1.00 | 0.42 | 0.95 | <0.001 |
| *φ ~ S* | **0.08** | **0.02** | **3.99** | **<0.001** | **<0.001** | -0.02 | 0.05 | -0.40 | 0.69 | 1.00 | <0.001 | 0.95 | 0.03 |
| *φ ~ R_0_* | **0.03** | **0.01** | **3.66** | **<0.001** | **<0.001** | -0.02 | 0.05 | -0.38 | 0.71 | 1.00 | <0.001 | 0.95 | 0.03 |
| *S ~ R_0_* | **0.15** | **0.02** | **7.68** | **<0.001** | **<0.001** | 0.16 | 0.13 | 1.24 | 0.21 | 1.00 | 0.09 | 0.95 | 0.12 |

**Table S12. Loadings of phylogenetically corrected principal component analysis (pPCA) for the aquatic (n=62 species) and the terrestrial taxa (n=477 species) for which full demographic data were available.** Headers of axes with associated eigenvalues >1, indicating retention to explain observed variation (Legendre & Legendre 2012), are shown in bold. Bold numbers indicate loading absolute values >0.50. Outputs correspond to the mean values of 10 imputed data sets.

| **Life history traits** | | **PC1** | **PC2** | PC3 | PC4 | PC5 | PC6 |
| --- | --- | --- | --- | --- | --- | --- | --- |
| **Turnover** | Generation time (*T*) | **0.81** | -0.08 | -0.24 | -0.36 | 0.17 | 0.34 |
| **Survival** | Rate of senescence (*H*) | **0.68** | 0.30 | -0.26 | 0.61 | -0.02 | 0.10 |
|  | Age at maturity (*L_α_*) | **0.76** | 0.06 | -0.45 | -0.22 | -0.18 | -0.37 |
| **Development** | Development (*γ*) | **-0.71** | -0.18 | -0.52 | 0.08 | 0.43 | -0.07 |
| **Reproduction** | Mean reproducitve output (*Φ*) | **-0.71** | 0.50 | -0.34 | -0.11 | -0.31 | 0.15 |
|  | Degree of iteroparity (*S*) | 0.12 | **0.92** | 0.19 | -0.12 | 0.29 | -0.09 |
| **Eigenvalue** | | **2.76** | **1.37** | 0.76 | 0.58 | 0.44 | 0.29 |
| **Proportion of variance explained** | | 44.65% | 22.09% | 12.21% | 9.34% | 7.10% | 4.61% |
| **Cumulative proportion of variance explained** | | 44.65% | 66.74% | 78.95% | 88.29% | 95.39% | 100.00% |
| **Pagel’s λ** | | 0.31 | | | | | |

**Table S13. Phylogenetically Generalised Least Squares (PGLS) regressions between life history traits for the aquatic (n=62 species) and the terrestrial taxa (n=477 species) for which full demographic data were available.** Here are presented the slope of the correlations between two life history traits (see Table S2 for interpretation of life history trait symbols), the effect of the realm, and the interaction with the second life history trait. *P*-adjusted is the adjusted *P*-value after a Bonferroni correction. Bold numbers represent significant correlations at *P*<0.05. Pagel’s *λ* quantifies phylogenetic signal (1: highest; 0: lowest), and its 95% confidence interval (CI). These results do not change qualitatively when phylogenetic imputations were carried out on the original dataset (Table S11).

| Traits | Slope | SD | *t*-value | *P* | *P*-adjusted | Realm | SD | *t*-value | *P* | *P*-adjusted | Pagel’s *λ* | CI | R^2^ |
| --- | --- | --- | --- | --- | --- | --- | --- | --- | --- | --- | --- | --- | --- |
| *T ~ H* | **1.69** | **0.24** | **6.95** | **<0.001** | **<0.001** | 0.89 | 1.10 | 0.81 | 0.42 | 1.00 | 0.78 | 0.95 | 0.15 |
| *T ~ Lα* | **0.86** | **0.07** | **13.06** | **<0.001** | **<0.001** | 0.12 | 0.29 | 0.43 | 0.67 | 1.00 | <0.001 | 0.95 | 0.38 |
| *T ~ γ* | **-5.26** | **0.95** | **-5.51** | **<0.001** | **<0.001** | 0.99 | 1.32 | 0.75 | 0.45 | 1.00 | 0.84 | 0.95 | 0.10 |
| *T ~ φ* | **-7.96** | **1.14** | **-7.00** | **<0.001** | **<0.001** | 0.97 | 1.37 | 0.70 | 0.48 | 1.00 | 0.86 | 0.95 | 0.15 |
| *T ~ S* | -0.01 | 0.11 | -0.07 | 0.95 | 0.95 | 1.10 | 1.79 | 0.62 | 0.54 | 1.00 | 0.91 | 0.95 | <0.001 |
| *H ~ Lα* | **0.16** | **0.02** | **9.55** | **<0.001** | **<0.001** | -0.02 | 0.07 | -0.24 | 0.81 | 1.00 | <0.001 | 0.95 | 0.24 |
| *H ~ γ* | **-1.38** | **0.21** | **-6.45** | **<0.001** | **<0.001** | 0.09 | 0.22 | 0.43 | 0.67 | 1.00 | 0.72 | 0.95 | 0.13 |
| *H ~ φ* | **-1.12** | **0.27** | **-4.12** | **<0.001** | **<0.001** | 0.10 | 0.26 | 0.41 | 0.69 | 1.00 | 0.78 | 0.95 | 0.06 |
| *H ~ S* | **0.08** | **0.03** | **3.19** | **<0.001** | **<0.001** | 0.11 | 0.30 | 0.39 | 0.70 | 1.00 | 0.83 | 0.95 | 0.03 |
| *Lα ~ γ* | **-3.68** | **0.64** | **-5.74** | **<0.001** | **<0.001** | 0.79 | 1.20 | 0.66 | 0.51 | 1.00 | 0.91 | 0.95 | 0.10 |
| *Lα ~ φ* | **-3.37** | **0.80** | **-4.19** | **<0.001** | **<0.001** | 0.80 | 1.28 | 0.63 | 0.53 | 1.00 | 0.92 | 0.95 | 0.06 |
| *Lα ~ S* | 0.04 | 0.08 | 0.47 | 0.64 | 0.66 | 0.84 | 1.54 | 0.55 | 0.58 | 1.00 | 0.94 | 0.95 | <0.001 |
| *γ ~ ρ* | **1.09** | **0.28** | **3.89** | **<0.001** | **<0.001** | -0.02 | 0.02 | -0.96 | 0.34 | 1.00 | <0.001 | 0.95 | 0.05 |
| *γ ~ φ* | **0.32** | **0.07** | **4.51** | **<0.001** | **<0.001** | -0.01 | 0.02 | -0.52 | 0.60 | 1.00 | <0.001 | 0.95 | 0.07 |
| *γ ~ S* | **-0.02** | **0.01** | **-3.30** | **<0.001** | **<0.001** | -0.01 | 0.02 | -0.54 | 0.59 | 1.00 | <0.001 | 0.95 | 0.04 |

**Table S14. Loadings of life history traits grouped into turnover, survival, development, and reproduction attributes, on the principal component axes for 512 sessile and 171 mobile species, separately.** Bold numbers indicate loading absolute values >50%. The first four principal component axes are represented. Eigenvalues >1 are shown in bold. Pagel’s *λ* describes the role of phylogenetic inertia in explaining the variation in the PCA, it ranges between 1 when life history trait differences are fully due to the phylogenetic structure as explained by Brownian motion, and 0 meaning no phylogenetic structuring in the pattern.

| **Life history traits** | | **Sessile** | | | | **Mobile** | | | |
| --- | --- | --- | --- | --- | --- | --- | --- | --- | --- |
|  |  | **PC1** | **PC2** | PC3 | PC4 | **PC1** | **PC2** | **PC3** | PC4 |
| **Turnover** | *T* | **0.85±0.00** | -0.08±0.01 | -0.08±0.02 | -0.26±0.07 | **0.79±0.00** | -0.25±0.02 | -0.44±0.01 | -0.10±0.03 |
| **Survival** | *H* | **0.72±0.01** | 0.21±0.01 | -0.30±0.08 | 0.42±0.10 | **0.73±0.01** | 0.36±0.03 | 0.22±0.03 | -0.48±0.03 |
|  | *L_α_* | **0.81±0.00** | 0.00±0.01 | -0.30±0.07 | -0.23±0.06 | **0.80±0.01** | -0.16±0.04 | -0.45±0.01 | 0.11±0.03 |
| **Development** | *γ* | **-0.72±0.01** | -0.18±0.01 | -0.45±0.11 | 0.00±0.02 | **-0.79±0.00** | 0.04±0.04 | -0.37±0.02 | -0.39±0.02 |
| **Reproduction** | *Φ* | **-0.67±0.01** | **0.54±0.01** | -0.23±0.05 | -0.15±0.05 | **-0.69±0.01** | 0.43±0.03 | -0.44±0.02 | 0.04±0.03 |
|  | *S* | 0.16±0.02 | **0.93±0.00** | 0.11±0.02 | -0.02±0.01 | 0.40±0.02 | **0.85±0.01** | -0.06±0.04 | 0.17±0.02 |
| **Eigenvalue** | | **3.02±0.03** | **1.35±0.01** | 0.75±0.01 | 0.53±0.01 | **3.99±0.07** | **1.60±0.04** | **1.04±0.02** | 0.62±0.02 |
| **Proportion of variance explained** | | 40.15±0.42% | 21.71±0.18% | 11.66±0.08% | 8.91±0.14% | 50.45±0.61% | 20.26±0.37% | 13.23±0.28% | 7.81±0.21% |
| **Pagel’s *λ*** | | 0.18±0.01 | | | | 0.36±0.01 | | | |

**Table S15. Loadings of phylogenetically corrected principal component analysis (pPCA) separated by kingdoms, with the life history traits grouped into turnover, survival, development, and reproduction attributes.** First are presented the results of the pPCA for the Animal kingdom (212 species), and then Plantae (463 species) and Chromista kingdoms (8 species) together. Bold numbers indicate loading absolute values >0.5. Eigenvalues >1 are in bold, indicating retention of the axis (Legendre & Legendre, 2012). Pagel’s *λ* describes the role of phylogenetic inertia in explaining the variation in the PCA, it ranges between 1 when life history trait differences are fully due to the phylogenetic structure as explained by Brownian motion, and 0 meaning no phylogenetic structuring in the pattern. The Plantae and Chromista kingdoms were merged due to the inability of the method to run only for Chromista due to its low sample size, but the pattern remains unaltered when analyzed only for plants (not shown). Here are presented the mean values and the standard errors of the values coming from the multiple imputed data sets.

|  | | **Animalia** | | | | **Plantae and Chromista** | | | |
| --- | --- | --- | --- | --- | --- | --- | --- | --- | --- |
|  |  | **PC1** | **PC2** | PC3 | PC4 | **PC1** | **PC2** | PC3 | PC4 |
| **Turnover** | *T* | **0.80±0.01** | -0.20±0.02 | -0.43±0.01 | -0.08±0.04 | **0.86±0.00** | -0.07±0.01 | -0.06±0.02 | -0.20±0.09 |
| **Survival** | *H* | **0.72±0.01** | 0.32±0.03 | 0.25±0.03 | -0.42±0.09 | **0.71±0.01** | 0.22±0.01 | -0.24±0.11 | 0.30±0.13 |
|  | *L_α_* | **0.80±0.01** | -0.08±0.04 | -0.47±0.01 | 0.08±0.04 | **0.81±0.00** | 0.01±0.01 | -0.22±0.09 | -0.18±0.08 |
| **Development** | *γ* | **-0.78±0.00** | 0.06±0.04 | -0.37±0.01 | -0.31±0.08 | **-0.72±0.01** | -0.20±0.01 | -0.34±0.15 | 0.00±0.02 |
| **Reproduction** | *Φ* | **-0.70±0.01** | 0.44±0.03 | -0.40±0.03 | 0.00±0.03 | **-0.67±0.01** | **0.55±0.01** | -0.16±0.07 | -0.15±0.06 |
|  | *S* | 0.37±0.02 | **0.87±0.01** | -0.02±0.04 | 0.15±0.03 | 0.15±0.02 | **0.92±0.00** | 0.07±0.04 | 0.00±0.02 |
| **Eigenvalue** | | **3.74±0.07** | **1.45±0.03** | 0.99±0.03 | 0.60±0.01 | **3.05±0.03** | **1.38±0.01** | 0.75±0.01 | 0.52±0.01 |
| **Proportion of variance explained** | | 50.49±0.53% | 19.65±0.30% | 13.34±0.35% | 8.13±0.19% | 47.63±0.41% | 21.49±0.11% | 11.78±0.17% | 8.13±0.13% |
| **Pagel’s *λ*** | | 0.31±0.02 | | | | 0.18±0.01 | | | |

**Table S16. Loadings of life history traits grouped into turnover, longevity, development and reproduction attributes, on the first four principal component axes for 638 terrestrial and 117 aquatic species separately.** Bold numbers indicate loading absolute values >50%. Eigenvalues >1 are showbn bold. Pagel’s *λ* describes the role of phylogenetic inertia in explaining the variation in the PCA, it ranges between 1 when life history trait differences are fully due to the phylogenetic structure as explained by Brownian motion, and 0 meaning no phylogenetic structuring in the pattern.

| **Life history traits** | | **Terrestrial** | | | | **Aquatic** | | | |
| --- | --- | --- | --- | --- | --- | --- | --- | --- | --- |
|  |  | **PC1** | **PC2** | PC3 | PC4 | **PC1** | **PC2** | PC3 | PC4 |
| **Turnover** | *T* | **0.85±0.00** | -0.07±0.01 | -0.05±0.04 | -0.20±0.09 | **0.75±0.01** | 0.04±0.05 | -0.18±0.17 | -0.14±0.05 |
| **Survival** | *H* | **0.71±0.01** | 0.24±0.01 | -0.15±0.11 | 0.32±0.15 | **0.77±0.01** | 0.18±0.07 | 0.12±0.08 | -0.27±0.11 |
|  | *L_α_* | **0.81±0.00** | 0.00±0.01 | -0.15±0.12 | -0.17±0.07 | **0.74±0.01** | 0.36±0.04 | -0.21±0.09 | 0.18±0.10 |
| **Development** | *γ* | **-0.72±0.01** | -0.19±0.01 | -0.21±0.16 | -0.01±0.02 | **-0.72±0.02** | 0.45±0.04 | -0.13±0.10 | -0.20±0.08 |
| **Reproduction** | *Φ* | **-0.68±0.01** | 0.53±0.01 | -0.14±0.10 | -0.11±0.04 | **-0.75±0.02** | **0.53±0.03** | -0.07±0.03 | 0.02±0.04 |
|  | *S* | 0.16±0.02 | **0.92±0.01** | 0.07±0.05 | -0.04±0.02 | 0.39±0.03 | **0.76±0.03** | 0.17±0.13 | 0.06±0.03 |
| **Eigenvalue** | | **3.08±0.03** | **1.39±0.01** | 0.75±0.01 | 0.55±0.01 | **3.18±0.07** | **1.45±0.07** | 0.82±0.04 | 0.52±0.03 |
| **Proportion of variance explained** | | 47.69±0.37% | 21.53±0.09% | 11.57±0.16% | 8.54±0.14% | 49.29±1.03% | 22.44±0.97% | 12.66±0.50% | 8.02±0.39% |
| **Pagel’s *λ*** | | 0.24±0.01 | | | | 0.19±0.02 | | | |

References

Bielby, J., Mace, G.M., Bininda-Emonds, O.R.P., Cardillo, M., Gittleman, J.L., Jones, K.E., Orme, C.D.L. & Purvis, A. (2007) The fast-slow continuum in mammalian life history: An empirical reevaluation. *The American naturalist*, **169**, 748–57.

Burns, J.H., Blomberg, S.P., Crone, E.E., Ehrlén, J., Knight, T.M., Pichancourt, J.-B., Ramula, S., Wardle, G.M. & Buckley, Y.M. (2010) Empirical tests of life-history evolution theory using phylogenetic analysis of plant demography. *Journal of Ecology*, **98**, 334–344.

Caswell, H. (2001) *Matrix Population Models: Construction, Analysis, and Interpretation*, 2nd edn. Sinauer Associates.

Chamberlain, S.A. & Szöcs, E. (2013) taxize: taxonomic search and retrieval in R. *F1000Research*, **2**, 191.

Demetrius, L. (1974) Demographic parameters and natural selection. *Proceedings of the National Academy of Sciences*, **71**, 4645–4647.

Gaillard, J.-M., Yoccoz, N.G., Lebreton, J.-D., Bonenfant, C., Devillard, S., Loison, A., Pontier, D. & Allaine, D. (2005) Generation time: A reliable metric to measure life-history variation among mammalian populations. *The American naturalist*, **166**, 119–123.

Grafen, A. (1989) The phylogenetic regression. *Phil. Trans. R. Soc. Lond. B*, **326**, 119–157.

Grosberg, R.K., Vermeij, G.J. & Wainwright, P.C. (2012) Biodiversity in water and on land. *Current Biology*, **22**, R900–R903.

Hinchliff, C.E., Smith, S.A., Allman, J.F., Burleigh, J.G., Chaudhary, R., Coghill, L.M., Crandall, K.A., Deng, J., Drew, B.T., Gazis, R., Gude, K., Hibbett, D.S., Katz, L.A., Laughinghouse, H.D., McTavish, E.J., Midford, P.E., Owen, C.L., Ree, R.H., Rees, J.A., Soltis, D.E., Williams, T. & Cranston, K.A. (2015) Synthesis of phylogeny and taxonomy into a comprehensive tree of life. *Proceedings of the National Academy of Sciences*, **112**, 12764–12769.

Honaker, J., King, G. & Blackwell, M. (2011) **Amelia** II: A Program for Missing Data. *Journal of Statistical Software*, **45**.

Keyfitz, N. (1977) What difference would it make if cancer were eradicated? An examination of the taeuber paradox. *Demography*, **14**, 411–418.

Legendre, P. & Legendre, L. (2012) *Numerical Ecology*, 3rd edn. Elsevier Science, Amsterdam.

Maddison, W. & Maddison, D. (2018) Mesquite: A modular system for evolutionary analysis.

Michonneau, F., Brown, J. & Winter, D. (2016) rotl: An R package to interact with the Open Tree of Life data. *Methods in Ecology and Evolution*, **7**, 1–17.

Midford, P.E., Garland, T.Jr. & Maddison, W. (2005) PDAP Package of Mesquite.

Morris, W. & Doak, D. (2002) *Quantitative Conservation Biology: Theory and Practice of Population Viability Analysis.*, Sinauer Associates, Sunderland, MA.

Paniw, M., Ozgul, A. & Salguero‐Gómez, R. (2018) Interactive life-history traits predict sensitivity of plants and animals to temporal autocorrelation. *Ecology Letters*, **21**, 275–286.

Paradis, E., Claude, J. & Strimmer, K. (2004) APE: Analyses of phylogenetics and evolution in R language. *Bioinformatics*, **20**, 289–290.

Revell, L.J. (2010) Phylogenetic signal and linear regression on species data. *Methods in Ecology and Evolution*, **1**, 319–329.

Revell, L.J. (2012) phytools: An R package for phylogenetic comparative biology (and other things). *Methods in Ecology and Evolution*, **3**, 217–223.

Salguero-Gómez, R., Jones, O.R., Jongejans, E., Blomberg, S.P., Hodgson, D.J., Mbeau-Ache, C., Zuidema, P.A., De Kroon, H. & Buckley, Y.M. (2016) Fast–slow continuum and reproductive strategies structure plant life-history variation worldwide. *Proceedings of the National Academy of Sciences*, **113**, 230–235.

Salguero-Gómez, R. & Plotkin, J.B. (2010) Matrix dimensions bias demographic inferences: implications for comparative plant demography. *The American Naturalist*, **176**, 710–722.

Silvertown, J. & Franco, M. (1993) Plant demography and habitat: A comparative approach. *Plant Species Biology*, **8**, 67–73.

Stearns, S.C. (1992) *The Evolution of Life Histories*, Oxford University Press, New York.

Tuljapurkar, S. & Haridas, C.V. (2006) Temporal autocorrelation and stochastic population growth. *Ecology Letters*, **9**, 327–337.
